# Semantic representations during language comprehension are affected by context

**DOI:** 10.1101/2021.12.15.472839

**Authors:** Fatma Deniz, Christine Tseng, Leila Wehbe, Tom Dupré la Tour, Jack L. Gallant

## Abstract

The meaning of words in natural language depends crucially on context. However, most neuroimaging studies of word meaning use isolated words and isolated sentences with little context. Because the brain may process natural language differently from how it processes simplified stimuli, there is a pressing need to determine whether prior results on word meaning generalize to natural language. fMRI was used to record human brain activity while four subjects (two female) read words in four conditions that vary in context: narratives, isolated sentences, blocks of semantically similar words, and isolated words. We then compared the signal-to-noise ratio (SNR) of evoked brain responses, and we used a voxelwise encoding modeling approach to compare the representation of semantic information across the four conditions. We find four consistent effects of varying context. First, stimuli with more context evoke brain responses with higher SNR across bilateral visual, temporal, parietal, and prefrontal cortices compared to stimuli with little context. Second, increasing context increases the representation of semantic information across bilateral temporal, parietal, and prefrontal cortices at the group level. In individual subjects, only natural language stimuli consistently evoke widespread representation of semantic information. Third, context affects voxel semantic tuning. Finally, models estimated using stimuli with little context do not generalize well to natural language. These results show that context has large effects on the quality of neuroimaging data and on the representation of meaning in the brain. Thus, neuroimaging studies that use stimuli with little context may not generalize well to the natural regime.

**Significance Statement:** Context is an important part of understanding the meaning of natural language, but most neuroimaging studies of meaning use isolated words and isolated sentences with little context. Here we examined whether the results of neuroimaging studies that use out-of-context stimuli generalize to natural language. We find that increasing context improves the quality of neuroimaging data and changes where and how semantic information is represented in the brain. These results suggest that findings from studies using out-of-context stimuli may not generalize to natural language used in daily life.

## Introduction

Language is our main means of communication and an integral part of daily life. Natural language comprehension requires extracting meaning from words that are embedded in context. However, most neuroimaging studies of word meaning use simplified stimuli consisting of isolated words or sentences (Price 2012). Natural language differs from isolated words and sentences in several ways. Natural language contains phonological and orthographic patterns, lexical semantics, syntactic structure, and compositional- and discourse-level semantics embedded in social context (Hagoort 2019). In contrast, isolated words and sentences only contain a few of these components (e.g., lexical meaning, local syntactic structure). (For concision, this paper will refer to all differences between natural language and isolated words/sentences as differences in “context.”)

Neuroimaging studies that use isolated words and sentences implicitly assume that their results will generalize to natural language. However, because the brain is a highly nonlinear dynamical system (Wu, David, and Gallant 2006; Breakspear 2017), the representation of semantic information may change depending on context (Poeppel et al. 2012; Hagoort 2019; Hamilton and Huth 2020). Indeed, contextual effects have been demonstrated clearly in other domains. For example, many neurons in the visual system respond differently to simplified stimuli compared to naturalistic stimuli (Simoncelli and Olshausen 2001; Ringach, Hawken, and Shapley 2002; David, Vinje, and Gallant 2004; Touryan, Felsen, and Dan 2005). However, few studies have examined whether insights about semantic representation from studies using simplified stimuli will generalize to natural language.

Results from past studies suggest that context has a large effect on semantic representation. Several natural language studies from our lab reported that semantic information is represented in a large, distributed network of brain regions including bilateral temporal, parietal, and prefrontal cortices, and that semantic information is represented independently of the presentation modality (Huth et al. 2016; Deniz et al. 2019). In contrast, studies that used isolated words or sentences as stimuli independently identified only a few brain regions that represent semantic information. These studies have separately identified angular gyrus, left inferior frontal gyrus (IFG), left ventromedial prefrontal cortex (vmPFC), left dorsolateral prefrontal cortex (dmPFC), anterior temporal lobe, lateral-, ventral-, and inferotemporal cortex, posterior cingulate gyrus, and posterior parietal cortex (for reviews see (Jeffrey R. Binder et al. 2009; Price 2010, 2012).

One way that context might affect neuroimaging results is by affecting the signal-to-noise ratio (SNR) of evoked brain responses (i.e., affecting the metabolic activity of the brain such that the repeatability of the recorded blood-oxygen-level-dependent (BOLD) response is affected). Although no language studies have explicitly looked at evoked BOLD SNR, several converging lines of evidence suggest that context does affect evoked SNR in language studies. (Lerner et al. 2011) examined how language context affects cross-subject correlations in brain responses, and they reported that as the amount of context increased, the number of voxels that were correlated across subjects also increased. These voxels were located in high-level brain regions including TPJ, precuneus, and mPFC. In contrast, voxel responses in sensory regions and the superior temporal sulcus (STS) were reliably correlated when stimuli with little context stimuli was presented to the subjects (also see (Hasson, Chen, and Honey 2015)). In addition, several contrast-based fMRI language studies reported that increasing context evoked larger and more widespread patterns of brain activity in posterior STS, TPJ, and mPFC (Mazoyer et al. 1993; Xu et al. 2005; Jobard et al. 2007). Finally, most subjects are more attentive when reading natural stories than when reading isolated words, and attention affects BOLD SNR (Bressler and Silver 2010).

Another more interesting way that context might affect neuroimaging results is by directly changing semantic representations in the brain (i.e., changing which voxels represent semantic information and/or the semantic tuning of those voxels). Context can change the way that subjects attend to semantic information, and semantic representations in many brain areas shift toward attended semantic categories (Çukur et al. 2013; Sprague, Saproo, and Serences 2015; Nastase et al. 2017). Context also changes the statistical structure of language stimuli, and these statistical changes can affect cognitive processes and representations in a variety of ways (Wu, David, and Gallant 2006; Dahmen et al. 2010; Breakspear 2017).

To test the hypotheses that context affects evoked SNR and semantic representations, we used fMRI and a voxelwise encoding model approach to directly compare four stimulus conditions that vary in context: Narratives, Sentences, Semantic Blocks, and Single Words (Figure 1). The Narratives condition consisted of four narrative stories used in our previous studies (Huth et al. 2016; Deniz et al. 2019; Popham et al. 2021). The other three conditions used sentences, blocks of semantically similar words, and individual words sampled from the narratives.

**Figure 1:**
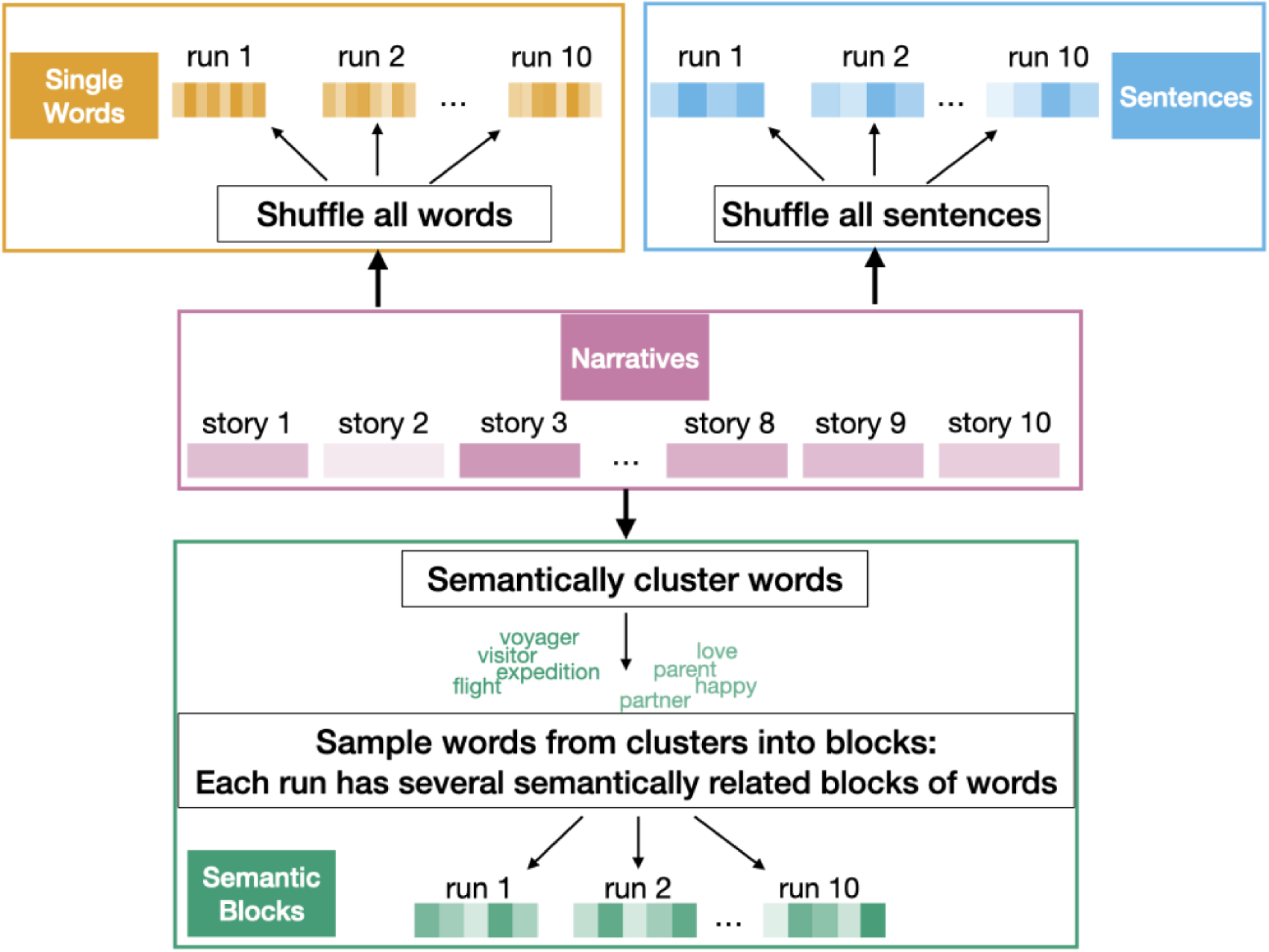
Stimulus conditions. The experiment contained four stimulus conditions that were based on the ten narratives used in Huth et al. (2016). The Single Words condition consisted of words sampled randomly from the ten narratives. The Semantic Blocks condition consisted of blocks of words sampled from clusters of semantically similar words from the ten narratives. There were 12 distinct clusters of semantically similar words, and blocks of words were created by randomly sampling 114 words from one word cluster for each block. The Sentences condition consisted of sentences sampled randomly from the ten narratives. Finally, the Narratives condition consisted of the ten original narratives.

## Materials and Methods

### Experimental Design and Statistical Analysis

#### Subjects

Functional data were collected from two males and two females: S1 (male, age 31), S2 (male, age 24), S3 (female, age 24), S4 (female, age 23). All subjects were healthy and had normal hearing, and normal or corrected-to-normal vision. All subjects were right handed according to the Edinburgh handedness inventory (Oldfield, 1971). Laterality scores were +70 (decile R.3) for S1, +95 (decile R.9) for S2, +90 (decile R.7) for S3, +80 (decile R.5) for S4.

#### MRI data collection

MRI data were collected on a 3T Siemens TIM Trio scanner with a 32-channel Siemens volume coil, located at the UC Berkeley Brain Imaging Center. Functional scans were collected using gradient echo EPI with repetition time (TR) = 2.0045s, echo time (TE) = 31ms, flip angle = 70 degrees, voxel size = 2.24 x 2.24 x 4.1 mm (slice thickness = 3.5 mm with 18% slice gap), matrix size = 100 x 100, and field of view = 224 x 224 mm. Thirty axial slices were prescribed to cover the entire cortex and were scanned in interleaved order. A custom-modified bipolar water excitation radiofrequency (RF) pulse was used to avoid signal from fat. Anatomical data were collected using a T1-weighted multi-echo MP-RAGE sequence on the same 3T scanner. Approximately 3.5 hours (214.85 minutes) of fMRI data was collected for each subject.

#### fMRI data pre-processing

The FMRIB Linear Image Registration Tool (FLIRT) from FSL 5.0 (M. Jenkinson and Smith 2001; Mark Jenkinson et al. 2002) was used to motion-correct each functional run. A high-quality template volume was then created for each run by averaging all volumes in the run across time. FLIRT was used to automatically align the template volume for each run to an overall template, which was chosen to be the temporal average of the first functional run for each subject. These automatic alignments were manually checked and adjusted as necessary to improve accuracy. The cross-run transformation matrix was then concatenated to the motion-correction transformation matrices obtained using MCFLIRT, and the concatenated transformation was used to resample the original data directly into the overall template space.

A 3rd order Savitsky-Golay filter with a 121-TR window was used to identify low-frequency voxel response drift. This drift was subtracted from the signal before further processing. Responses for each run were z-scored separately before voxelwise modeling. In addition, 10 TRs were discarded from the beginning and the end (20 TRs total) of each run.

#### Cortical surface reconstruction and visualization

Freesurfer (Dale, Fischl, and Sereno 1999) was used to generate cortical surface meshes from the T1-weighted anatomical scans. Before surface reconstruction, Blender and pycortex (http://pycortex.org; (Gao et al. 2015)) were used to carefully hand-check and correct anatomical surface segmentations. To aid in cortical flattening, Blender and pycortex were used to remove the surface crossing the corpus callosum and relaxation cuts were made into the surface of each hemisphere. The calcarine sulcus cut was made at the horizontal meridian in V1 as identified from retinotopic mapping data.

Pycortex (Gao et al. 2015) was used to align functional images to the cortical surface. The line-nearest scheme in pycortex was used to project functional data onto the surface for visualization and subsequent analysis. The line-nearest scheme samples the functional data at 64 evenly-spaced intervals between the inner (white matter) and outer (pial) surfaces of the cortex and averages the samples. Samples are taken using nearest-neighbor interpolation, in which each sample is given the value of its enclosing voxel.

#### Stimuli

Stimuli for all four conditions were generated from ten spoken stories from The Moth Radio Hour (used previously in (Huth et al. 2016)). In each story, a speaker tells an autobiographical story in front of a live audience. The ten selected stories are 10-15 min long, cover a wide range of topics, and are highly engaging. Transcriptions of these stories were used to generate the stimuli.

#### Story transcription

Each story was manually transcribed by one listener, and this transcription was checked by a second listener. Certain sounds (e.g., laughter, lip-smacking, and breathing) were also transcribed in order to improve the accuracy of the automated alignment. The audio of each story was downsampled to 11.5 kHz and the Penn Phonetics Lab Forced Aligner (P2FA; (Yuan and Liberman 2008)) was used to automatically align the audio to the transcript. P2FA uses a phonetic hidden Markov model to find the temporal onset and offset of each word and phoneme. The Carnegie Mellon University pronouncing dictionary was used to guess the pronunciation of each word. The Arpabet phonetic notation was used when necessary to manually add words and word fragments that appeared in the transcript but not in the pronouncing dictionary.

After automatic alignment was complete, Praat (Boersman and Weenink 2014) was used to manually check and correct each aligned transcript. The corrected, aligned transcript was then spot-checked for accuracy by a different listener. Finally, Praat’s TextGrid object was used to convert the aligned transcripts into word representations. The word representation of each story is a list of pairs (W, t), where W is a word and t is the time in seconds.

#### Stimulus Conditions

To evaluate the effect of context on evoked SNR and semantic representation in the brain, four stimulus conditions with different amounts of context were created. These four conditions were Narratives, Sentences, Semantic Blocks, and Single Words.

The Narratives condition consisted of four narratives from The Moth Radio Hour (“undertheinfluence”, “souls”, “life”, “wheretheressmoke”). These four narratives were chosen from the ten narratives used in (Huth et al. 2016). Each narrative was presented in a separate ∼10-minute scanning run. One narrative (“wheretheressmoke”) was used as the model validation stimulus, and it was presented twice for each subject.

The Sentences condition consisted of sentences randomly sampled from the ten narratives used in (Huth et al. 2016). Sentence boundaries were marked manually, resulting in 1450 sentences with a median sentence length of 13 words (min=5 words, max=40 words). Sentences were presented in four unique ∼10-minute scanning runs. One run was used as the model validation stimulus, and it was presented twice for each subject.

The Semantic Blocks condition consisted of blocks of semantically clustered words from the ten narratives used in (Huth et al. 2016). The motivation for this condition was to mimic the timescale on which semantic topics change in natural language without including grammatical and syntactic components. The semantic word clusters were designed to elicit maximally different voxel responses. To create the clusters, each word was first transformed into its semantic model representation (see Voxelwise model fitting below). The semantic model representation for each word was then projected onto the first ten principal components of the semantic model weights estimated in (Huth et al. 2016). Finally, the projections were clustered with k-means clustering (k=12) to create 12 word clusters. During each scanning run, subjects saw 12 different blocks of 114 words each. The words in each block were sampled from one of the word clusters, and eight different word clusters were sampled in each run. The frequency with which each cluster was sampled was matched to the frequency with which words from that cluster appeared in the ten narratives. Blocks were presented in four unique ∼10-minute long runs. One run was used as the model validation stimulus, and it was presented twice for each subject.

The Single Words condition consisted of words randomly sampled without replacement from the ten narratives used in (Huth et al. 2016). There were 21743 appearances of 2868 unique words across the narratives, and each appearance was sampled uniformly. Words were presented in four unique 10-minute scanning runs. One run was used as the model validation stimulus, and it was presented twice for each subject.

For the Sentences, Semantic Blocks, and Single Words conditions, text descriptions of auditory sounds (e.g., laughter and applause) in the ten narratives were removed. In addition, obvious transcription errors were removed from the list of narrative words for the Semantic Blocks and Single Words conditions. Words that did not make sense by themselves (e.g., “tai”, “chi”) were also removed. There were five such words: “tai”, “chi”, “deja”, “vu”, and “sub.”

#### Stimulus presentation

In all conditions, words were presented individually at the center of the screen using Rapid Serial Visual Presentation (RSVP) (Forster 1970; Buchweitz et al. 2009). Words in the Narratives and Sentences conditions were presented with the same timing and duration as in the original spoken stories. Words in the Semantic Blocks and Single Words conditions were presented for a baseline of 400 ms with an additional 10 ms for every character. For example, the word “apple” would be presented for 400 ms + 10 ms/character * (5 characters) = 450 ms. The word presentation timing was determined after extensive pilot testing before the experiment was run. The resulting parameters provided a good balance between readability and keeping subject engagement.

For subjects S1, S2, and S4, the four conditions were presented in 15 runs over two scanning sessions. Each condition was presented in a separate run, and the runs were interleaved in each session. In the first session, the conditions were presented in the order: Single Words, Semantic Blocks (validation stimulus), Sentences, Single Words (validation stimulus), Semantic Blocks, Sentences (validation stimulus), Semantic Blocks, Sentences. In the second session, the conditions were presented in the order: Sentences, Single Words (validation stimulus), Semantic Blocks, Single Words, Semantic Blocks (validation stimulus), Single Words, Sentences (validation stimulus). Conditions were presented in the same order for subjects S1, S2, and S4. For subject S3, the four conditions were presented in four scanning sessions. Each condition was presented in a separate scanning session, and each session contained 8 runs (including two repetitions of the validation stimulus). The stimuli used for this paper was a subset of the total stimuli presented in the four sessions. Although the stimuli were presented differently for subject S3, the results for subject S3 are consistent with the other three subjects.

The pygame library in Python was used to display black text on a gray background at 34 horizontal and 27 vertical degrees of visual angle. Letters were presented at average 6 (min=1, max=16) horizontal and 3 vertical degrees of visual angle. A white fixation cross was present at the center of the display. Subjects were asked to fixate while reading the text. Eye movements were monitored at 60 Hz throughout the scanning sessions using a custom-built camera system equipped with an infrared source (Avotec) and the ViewPoint EyeTracker software suite (Arrington Research). The eye tracker was calibrated before each session of data acquisition.

#### Explainable variance (EV)

To measure the functional SNR of each stimulus condition, we computed the explainable variance (EV). EV was computed as the amount of variance in the response of a voxel that can be explained by the mean response of the voxel across multiple repetitions of the same stimulus. Formally, if the responses of a voxel to a repeated stimulus is expressed as a matrix Y with dimensions (# of TRs in each repetition, # of stimulus repetitions), then EV is given by

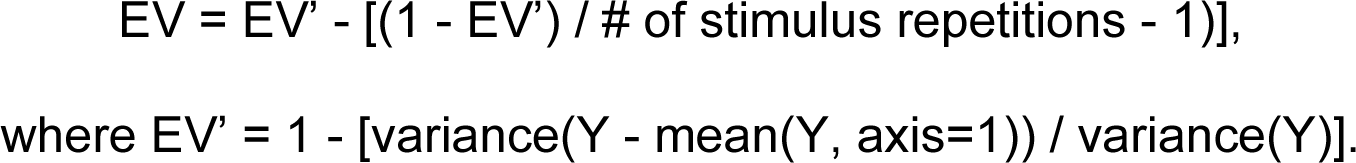

Note that this is the same as the coefficient of determination (R2) where the model prediction is the mean response across stimulus repetitions. For each condition, EV was computed from the two repeated validation runs.

#### Voxelwise model fitting and validation

To identify voxels that represent semantic information, a linearized encoding model (Nishimoto et al. 2011; Huth et al. 2012, 2016) was fit to every cortical voxel in each subject’s brain. The linearized encoding model consisted of one feature space designed to represent semantic information in the stimuli (the semantic feature space), and four feature spaces designed to represent low-level linguistic information in the stimuli. In the semantic feature space, the semantic content of each word was represented by the word’s co-occurrence statistics with the 985 words in Wikipedia’s List of 1000 basic words (Huth et al., 2016). Thus, each word was represented by a 985-long vector in the semantic feature space. The co-occurrence statistics were computed over a large text corpus that included the ten narrative stories used in Huth et al. (2016), several books from Project Gutenberg, a wide variety of Wikipedia pages, and a broad selection of reddit.com user comments (Huth et al. 2016). The four low-level feature spaces were word rate (1 parameter), number of letters (1 parameter), letters (26 parameters), and word length variation per TR (1 parameter). Together, the five feature spaces had 1014 features.

The features passed through three additional preprocessing steps before being fit to BOLD responses. First, to account for the hemodynamic response, a separate finite impulse response (FIR) filter with four delays was fit for each of the 1014 features, resulting in 4056 final features. This was accomplished by concatenating copies of the features delayed by 1, 2, 3, and 4 TRs (approximately 2, 4, 6, and 8 seconds). Taking the dot product of this concatenated feature space with a set of linear weights is functionally equivalent to convolving the undelayed features with a linear temporal kernel that has non-zero entries for 1-, 2-, 3-, and 4-time point delays. Second, 10 TRs were discarded from the beginning and the end (20 TRs total) of each run. Third, each feature was z-scored separately within each run. This was done so that the features would be on the same scale as the BOLD responses, which were also z-scored within each run.

A single joint model consisting of the 4056 features were fit to BOLD responses using banded ridge regression (Nunez-Elizalde, Huth, and Gallant 2019) and the himalaya Python package ((Dupré la Tour et al. 2022), see Code Accessibility). A separate model was fit for every voxel in every subject and condition. For every model, a regularization parameter was estimated for each of the five feature spaces using a random search. In the random search, 1000 normalized hyperparameter candidates were sampled from a Dirichlet distribution and scaled by 30 log-spaced values ranging from 10^-5 to 10^20. The best normalized hyperparameter candidate and scaling were selected for each feature space for each voxel. Finally, models were fit again on the BOLD responses with the selected hyperparameters.

To validate the models, estimated feature weights were used to predict responses to a separate, held-out validation dataset. Validation stimuli for the Narratives condition consisted of two repeated presentations of the narrative “wheretheressmoke” (Huth et al. 2016). Validation stimuli for the Sentences, Semantic Blocks, and Single Words conditions consisted of two repeated presentations of one run for each condition. Prediction accuracy was then computed by estimating the contribution of each feature space to the total prediction accuracy of the joint voxelwise model using the “correlation_score_split” function in the himalaya Python package (see also (St-Yves and Naselaris 2018), “Feature map contribution to the prediction accuracy”). This function computes the correlation between the predicted BOLD response from one feature space and the average BOLD response across the two validation runs, while accounting for the magnitude of the predictions from each feature space with respect to the other feature spaces in the joint model. The contribution from the semantic feature space is shown as semantic model prediction accuracy in Figures 4 and 5.

Statistical significance for each condition was computed with permutation testing. A null distribution was generated by permuting 10-TR blocks of the average validation BOLD response 5000 times and computing the prediction accuracy for each permutation (10 TRs were blocked to account for temporal autocorrelations in the BOLD signal). Resulting p values were corrected for multiple comparisons within each subject using the false discovery rate (FDR) procedure (Benjamini and Hochberg 1995).

#### Tuning shifts

To determine how semantic tuning changes between the Sentences and Narratives conditions, we looked at the difference between the estimated semantic model weights in the two conditions. First, temporal information was removed from the semantic model weights by averaging across the four delays for each semantic feature. Semantic model weights were then normalized by their L2-norm for each voxel, subject, and condition separately. This was done to ensure that the semantic model weights in both conditions are on the same numerical scale. Finally, the normalized semantic model weights estimated in the Sentences condition were subtracted from the normalized semantic model weights estimated in the Narratives condition.

To interpret the resulting difference vectors, we used principal components analysis (PCA) to recover a low-dimensional subspace. The difference vector for each voxel in each subject was scaled by the voxel’s minimum semantic model prediction accuracy between the Sentences and Narratives conditions. This was done to avoid including noise from voxels that were poorly predicted in either condition. We then applied PCA to the scaled difference vectors, yielding 985 PCs per subject. Partial scree plots showing the proportion of variance explained by the PCs in each subject are shown in Extended Data Figure 8-1. We projected each subject’s difference vectors onto the first three PCs for interpretation and visualization.

#### Cross-condition voxelwise model fitting

Estimated semantic model weights from the Single Words, Semantic Blocks and Sentences conditions were used to predict voxel responses in the Narratives condition. Prediction accuracy was computed as Pearson’s correlation coefficient between the predicted BOLD response using semantic model weights from the Single Words, Semantic Blocks, or Sentences condition and the average BOLD response across the two validation runs in the Narratives condition. In addition, estimated semantic model weights from the Single Words and Semantic Blocks conditions were used to predict voxel responses in the Sentences condition. Prediction accuracy was computed as Pearson’s correlation coefficient between the predicted BOLD response using semantic model weights from the Single Words or Semantic Blocks condition and the average BOLD response across the two validation runs in the Sentences condition.

All model fitting and analysis was performed using custom software written in Python, making heavy use of NumPy (Harris et al. 2020) and SciPy (Virtanen et al. 2020). Analysis and visualizations were developed using iPython (Perez and Granger 2007), the interactive programming and visualization environment jupyter notebook (Kluyver et al. 2016), Pycortex (Gao et al. 2015), and Matplotlib (Hunter 2007).

#### Code Accessibility

The himalaya package is publicly available on GitHub (https://github.com/gallantlab/himalaya).

## Results

The goal of this study was to understand whether context affects evoked SNR and whether it affects semantic representations in the brain. Previous studies suggest that both evoked SNR and semantic representations will differ across the four experimental conditions (Single Words, Semantic Blocks, Sentences, and Narratives). Here, we analyzed evoked SNR and semantic representations for each of the four conditions in individual subjects.

To estimate evoked SNR, we computed the reliability of voxel responses across repetitions of the same stimulus. Several different sources of noise can influence the variability of voxel responses across stimulus repetitions: magnetic inhomogeneity, voxel response variability, and variability in subject attention or vigilance. Because these sources are independent across stimulus repetitions, pooling voxel responses across repetitions averages out the noise and provides a good estimate of the evoked SNR. In this study, we used explainable variance (EV) as a measure of reliability and computed the EV for two repetitions of one run in each condition to estimate evoked SNR (see Methods).

Figure 3 shows EV for the four conditions in one typical subject (S1) (see Extended Data Figure 3-1 for voxels with significant EV; see Extended Data Figure 3-2 for unthresholded EV for subjects 2-4). In the Single Words condition, appreciable EV is only found in a few scattered voxels located in bilateral primary visual cortex, STS, and inferior frontal gyrus (IFG) (Figure 3a). The number of voxels with significant EV (p<0.05, FDR-corrected) in the Single Words condition is 256, 1198, 0, and 0 for subjects 1-4, respectively. A similar pattern is seen in the Semantic Blocks condition, where appreciable EV is only found in a few scattered voxels located in bilateral primary visual cortex, STS, and IFG (Figure 3b). The number of voxels with significant EV (p<0.05, FDR-corrected) in the Semantic Blocks condition is 324, 1613, 1201, and 0 for subjects 1-4, respectively. In contrast, both the Sentences and Narratives conditions produce high EV in many voxels located in bilateral visual, parietal, temporal, and prefrontal cortices (Figures 3c and 3d). The number of voxels with significant EV (p<0.05, FDR-corrected) in the Sentences condition is 4225, 11697, 2359, and 7251 for subjects 1-4, respectively. The number of voxels with significant EV (p<0.05, FDR-corrected) in the Narratives condition is 7622, 8062, 7059, and 2931 for subjects 1-4, respectively. Together, these results show that increasing context increases evoked SNR in bilateral visual, temporal, parietal, and prefrontal cortices.

To quantify semantic representation, we used a voxelwise encoding model (VM) procedure and a semantic feature space to identify voxels that represent semantic information in each condition (Figure 2). We first extracted semantic features and four types of low-level linguistic features from the stimulus words in each condition separately (see Methods). We then used banded ridge regression (Nunez-Elizalde, Huth, and Gallant 2019) to fit a joint encoding model for each voxel, subject, and condition. Finally, we split the joint model prediction accuracy across the five feature spaces to estimate the prediction accuracy for each feature space. Here we refer to voxels that had a significant semantic model prediction accuracy (see Methods) as “semantically selective voxels.”

**Figure 2:**
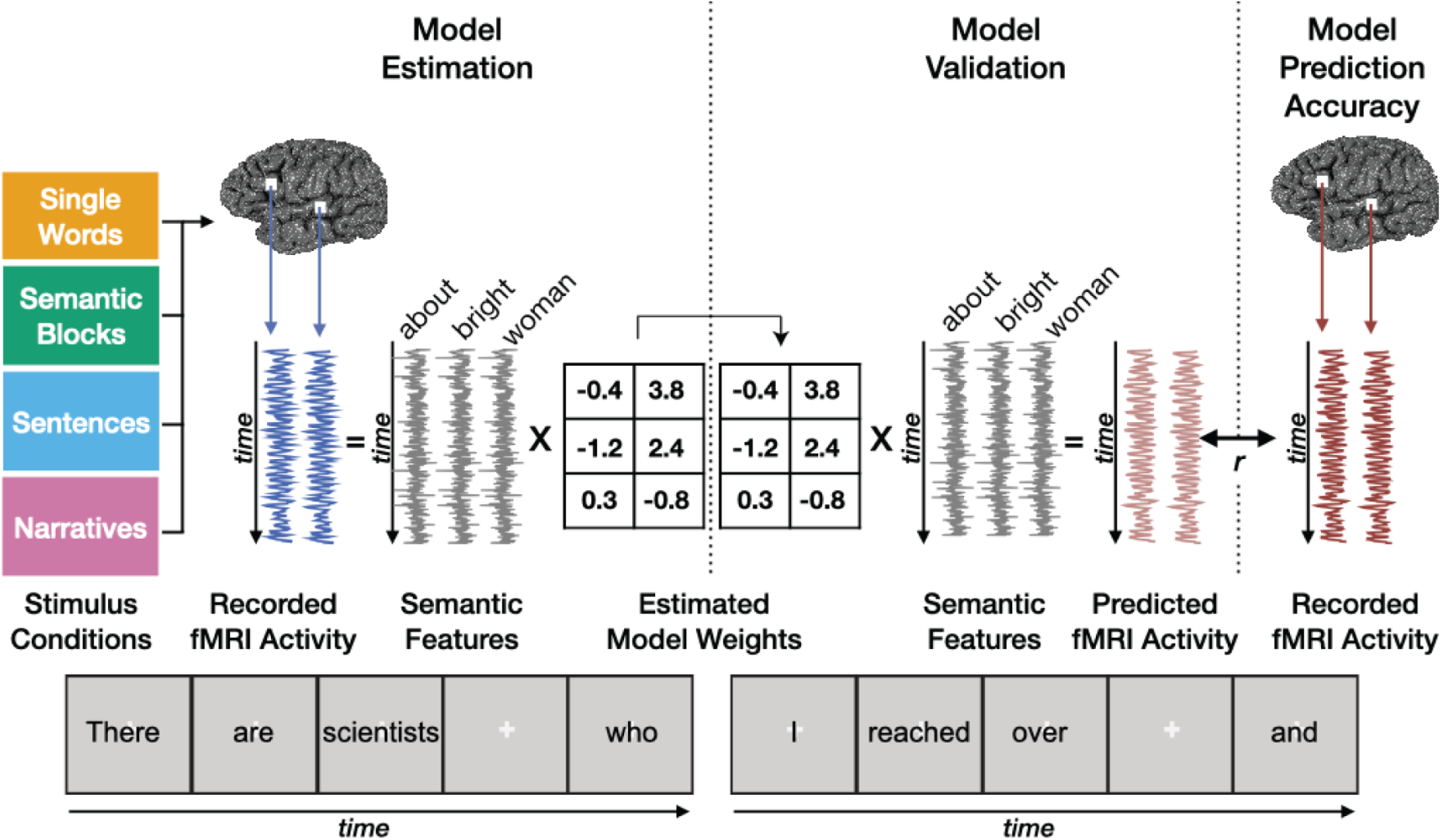
Voxelwise Modeling. Four subjects read words from the four stimulus conditions while BOLD responses were recorded. Each stimulus word was projected into a 985-dimensional word embedding space that was independently constructed using word co-occurrence statistics from a large corpus (Semantic Features). A finite impulse response (FIR) regularized regression model was estimated separately for each voxel in every subject and condition using banded ridge regression (Nunez-Elizalde et al. 2019). The estimated model weights were then used to predict BOLD responses to a separate, held-out validation stimulus. Model prediction accuracy was quantified as the correlation (r) between the predicted and recorded BOLD responses to the validation stimulus.

Figure 4 shows semantic model prediction accuracy for semantically selective voxels for the four conditions in one typical subject (S1) (see Extended Data Figure 4-1 for additional subjects; see Extended Data Figure 4-2 for unthresholded semantic model prediction accuracy for all subjects). In the Single Words condition, no voxels are semantically selective in any of the four subjects (Figure 4a and Extended Data Figure 4-3, p<0.05, FDR corrected). In the Semantic Blocks condition, scattered voxels along the left STS and left IFG are semantically selective (Figure 4b, p<0.05, FDR corrected). The number of semantically selective voxels (p<0.05, FDR corrected) in the Semantic Blocks condition is 708, 0, 0, and 0 for subjects 1-4, respectively (Extended Data Figure 4-3). In the Sentences condition, voxels in the left angular gyrus, left STG, bilateral STS, bilateral ventral precuneus, bilateral ventral premotor speech area (sPMv), bilateral superior frontal sulcus (SFS), and left superior frontal gyrus (SFG) are semantically selective (Figure 4c, p<0.05, FDR corrected). The number of semantically selective voxels (p<0.05, FDR-corrected) in the Sentences condition is 1566, 2581, 0, and 0 for subjects 1-4, respectively (Extended Data Figure 4-3). Finally, in the Narratives condition, voxels in bilateral angular gyrus, bilateral STS, bilateral STG, bilateral temporal parietal junction (TPJ), bilateral sPMv, bilateral ventral precuneus, bilateral SFS, bilateral SFG, bilateral IFG, left inferior parietal lobule (IPL), and left posterior cingulate gyrus are semantically selective (Figure 4d, p<0.05, FDR corrected). The number of semantically selective voxels (p<0.05, FDR-corrected) in the Narratives condition is 4745, 7355, 7786, and 1757 for subjects 1-4, respectively (Extended Data Figure 4-3). Together, these results suggest that increasing context increases the representation of semantic information in bilateral temporal, parietal, and prefrontal cortices. These results also suggest that this effect is highly variable in individual subjects for non-natural language stimuli (Semantic Blocks, Sentences) but not for natural language stimuli (Narratives).

**Figure 3.**
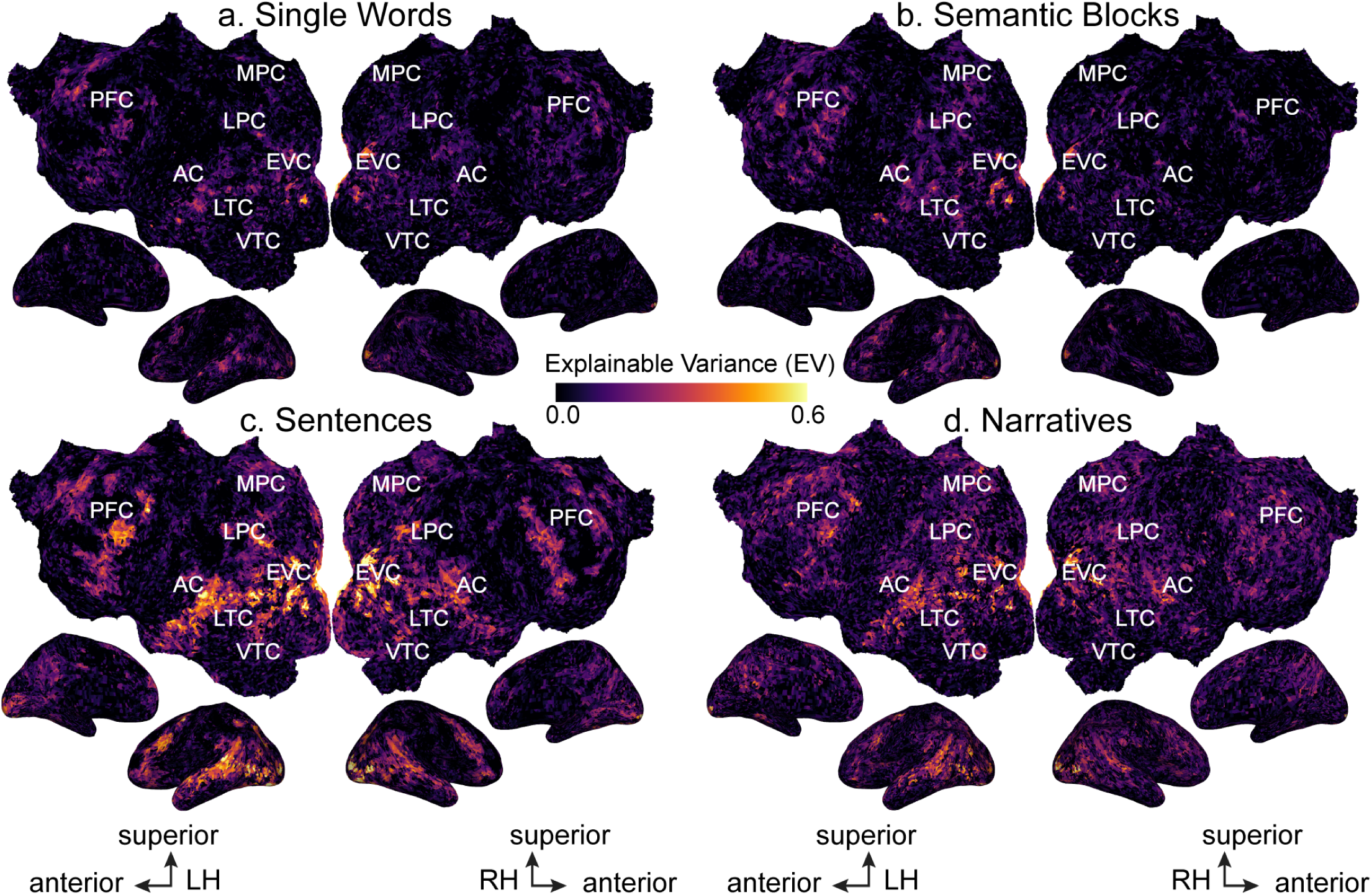
Explainable variance (EV) for the four conditions across the cortical surface. EV for the four conditions is shown for one subject (S1) on the subject’s flattened cortical surface. EV was computed as an estimate of the evoked signal-to-noise ratio (SNR). Here EV is given by the color scale shown in the middle, and voxels that have high EV (i.e., high evoked SNR) appear yellow. (LH: Left Hemisphere, RH: Right Hemisphere, AC: auditory cortex, EVC: early visual cortex, LTC: lateral temporal cortex, VTC: ventral temporal cortex, LPC: lateral parietal cortex, MPC: medial parietal cortex, PFC: prefrontal cortex) The format is the same in all panels. **a.** EV was computed for the Single Words condition and is shown on the flattened cortical surface of subject S1. Scattered voxels in bilateral primary visual cortex, superior temporal sulcus (STS), and inferior frontal gyrus (IFG) have high EV. **b.** EV was computed for the Semantic Blocks condition. Similar to the Single Words condition, scattered voxels in bilateral primary visual cortex, STS, and IFG have high EV. **c.** EV was computed for the Sentences condition. Many voxels in bilateral visual, parietal, temporal, and prefrontal cortices have high EV. **d.** EV was computed for the Narratives condition. Similar to the Sentences condition, voxels in bilateral visual, parietal, temporal, and prefrontal cortices have high EV. Together, these results show that increasing context increases evoked SNR in bilateral visual, temporal, parietal, and prefrontal cortices. (See Extended Data Figure 3-1 for significant EV voxels for subject S1 and Extended Data Figure 3-2 for EV for all subjects.)

**Figure 4.**
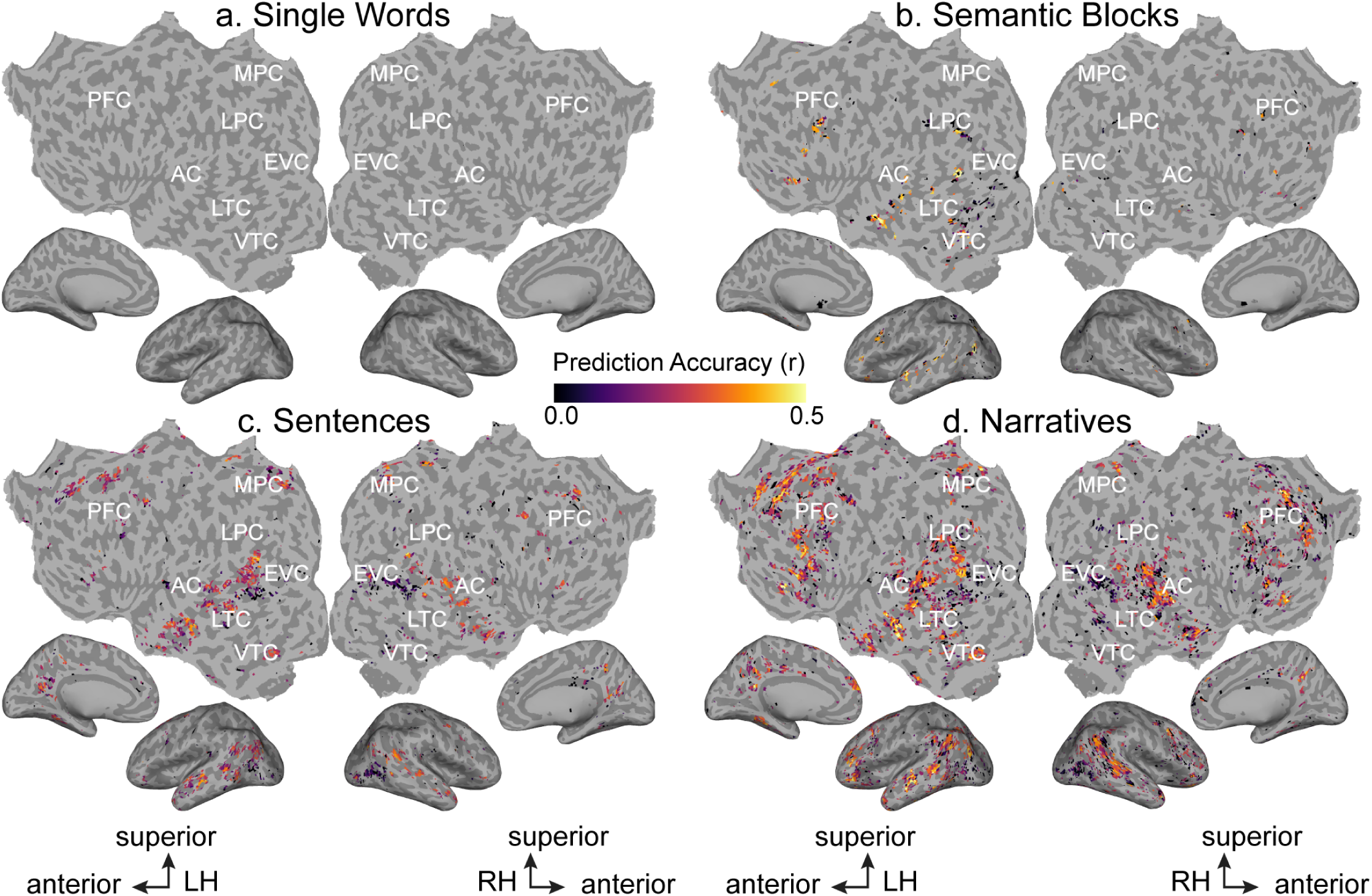
Semantic model prediction accuracy for the four conditions across the cortical surface. Semantic model prediction accuracy in the four conditions is shown on the flattened cortical surface of one subject (S1; see Extended Data Figure 4-1 and 4-2 for all subjects). Voxelwise modeling was first used to estimate semantic model weights in the four conditions. Semantic model prediction accuracy was then computed as the correlation (r) between the subject’s recorded BOLD activity to the held-out validation stimulus and the BOLD activity predicted by the semantic model. In each panel, only voxels with significant semantic model prediction accuracy (p<0.05, FDR corrected) are shown. Prediction accuracy is given by the color scale in the middle, and voxels that have a high prediction accuracy appear yellow. Voxels for which the semantic model prediction accuracy is not statistically significant are shown in gray. (LH: Left Hemisphere, RH: Right Hemisphere, AC: auditory cortex, EVC: early visual cortex, LTC: lateral temporal cortex, VTC: ventral temporal cortex, LPC: lateral parietal cortex, MPC: medial parietal cortex, PFC: prefrontal cortex) **a.** Semantic model prediction accuracy was computed for the Single Words condition. No voxels are significantly predicted in the Single Words condition (see Extended Data Figure 4-3 for the number of semantically selective voxels for the four conditions for all subjects). **b.** Semantic model prediction accuracy was computed for the Semantic Blocks condition. The format is the same as panel **a**. Voxels in left STS and IFG are significantly predicted. **c.** Semantic model prediction accuracy was computed for the Sentences condition. The format is the same as panel **a**. Voxels in left angular gyrus, left STG, bilateral STS, bilateral ventral precuneus, bilateral ventral premotor speech area (sPMv), bilateral superior frontal sulcus (SFS), and left superior frontal gyrus (SFG) are significantly predicted. **d.** Semantic model prediction accuracy was computed for the Narratives condition. The format is the same as panel **a.** Voxels in bilateral angular gyrus, bilateral STS, bilateral STG, bilateral temporal parietal junction (TPJ), bilateral sPMv, bilateral ventral precuneus, bilateral SFS, bilateral SFG, bilateral IFG, left inferior parietal lobule (IPL), and left posterior cingulate gyrus are significantly predicted. Together, these results suggest that increasing context increases the representation of semantic information in bilateral temporal, parietal, and prefrontal cortices.

The results presented in Figure 4 were obtained in each subject’s native brain space. To determine how the representation of semantic information varies across subjects for the four conditions, we transformed the semantic encoding model results obtained for each subject into the standard MNI brain space (Deniz et al. 2019). Figure 5 shows the mean unthresholded model prediction accuracy across subjects (Figure 5a-d) and the number of subjects for which each voxel is semantically selective (Figure 5e-h) for each condition. In the Single Words condition, no voxels are semantically selective in any of the four subjects (Figure 5a and 5e, p<0.05, FDR corrected). In the Semantic Blocks condition, scattered voxels in left STS are semantically selective in two out of four subjects (Figure 5b and 5f, p<0.05, FDR corrected). In the Sentences condition, voxels in the bilateral STS, left STG, bilateral ventral precuneus, bilateral angular gyrus, bilateral SFS, and bilateral premotor cortex are semantically selective in two out of four subjects (Figure 5c and 5g, p<0.05, FDR corrected). Finally, in the Narratives condition, voxels in bilateral angular gyrus, bilateral STS, right STG, right anterior temporal lobe, bilateral SFS and SFG, left IFG, left IPL, bilateral ventral precuneus, and bilateral posterior cingulate gyrus are semantically selective in all subjects (Figure 5d and 5h, p<0.05, FDR corrected), and voxels in left STG and right IFG are semantically selective in three out of four subjects (Figure 5d and 5h, p<0.05, FDR corrected). These results are consistent with those in Figure 4, and they suggest that increasing stimulus context increases the representation of semantic information across the cortical surface at the group level. In addition, this effect is inconsistent across individual subjects for non-natural stimuli (Semantic Blocks, Sentences) but not natural stimuli (Narratives).

**Figure 5.**
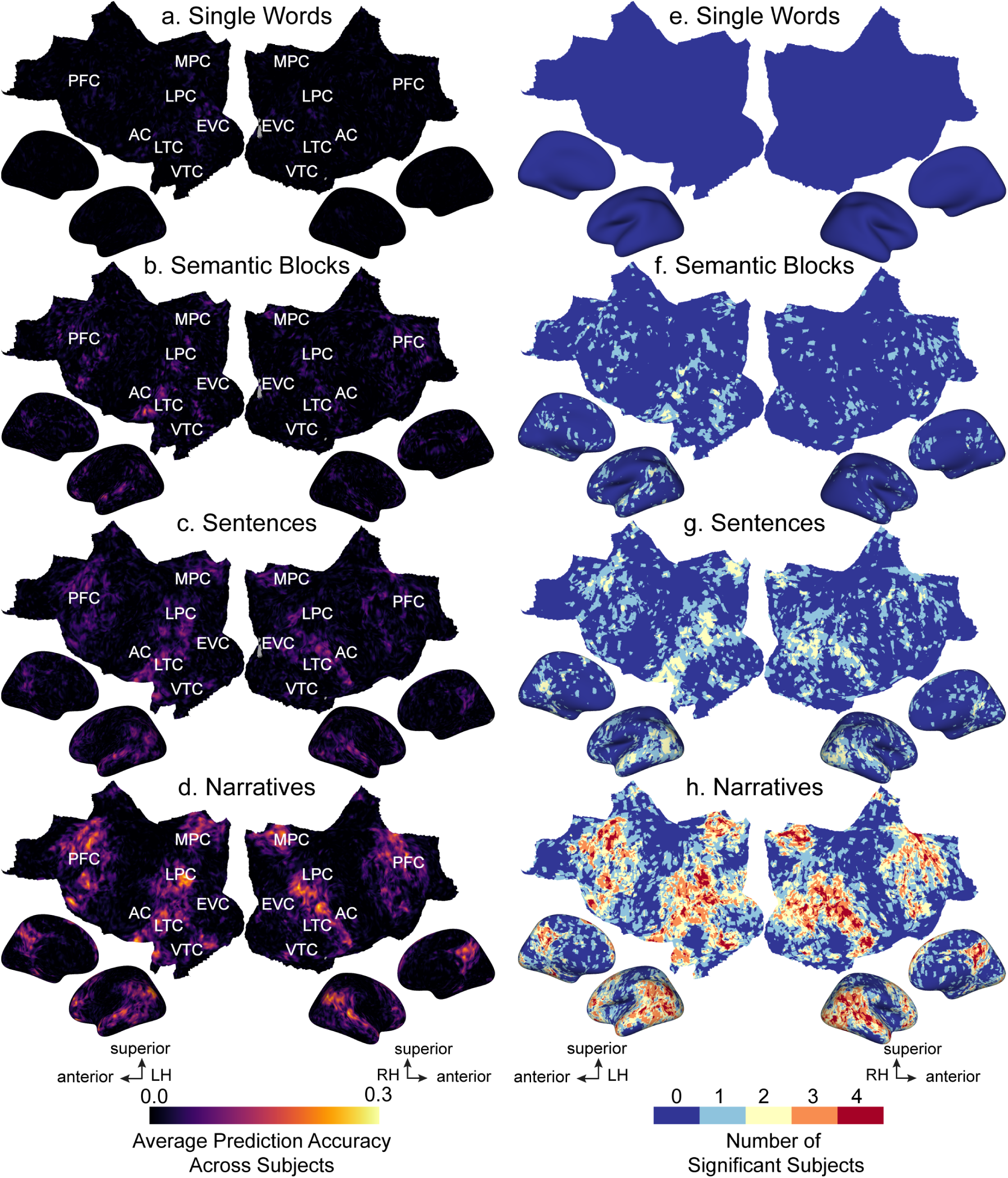
Semantic model prediction accuracy across all subjects for the four conditions in standard brain space. Semantic model prediction accuracy was first computed for each subject and for each condition as described in Figure 4. These individualized predictions were then projected into the standard MNI brain space. **a.-d.** Average prediction accuracy across the four subjects is computed for each MNI voxel and shown for each condition on the cortical surface of the MNI brain. Average prediction accuracy is given by the color scale, and voxels with higher prediction accuracy appear brighter. (LH: Left Hemisphere, RH: Right Hemisphere, AC: auditory cortex, EVC: early visual cortex, LTC: lateral temporal cortex, VTC: ventral temporal cortex, LPC: lateral parietal cortex, MPC: medial parietal cortex, PFC: prefrontal cortex) **a.** In the Single Words condition, average prediction accuracy is low across the cortical surface. **b.** In the Semantic Blocks condition, average prediction accuracy is high in voxels in left anterior STS. **c.** In the Sentences condition, average prediction accuracy is high in bilateral STS, STG, anterior temporal lobe, angular gyrus, ventral precuneus, SFS, and SFG. **d.** In the Narratives condition, average prediction accuracy is very high in bilateral STS, STG, MTG, anterior temporal lobe, angular gyrus, IPL, ventral precuneus, posterior cingulate gyrus, Broca’s area, IFG, SFS, SFG, and left posterior inferior temporal sulcus. **e.-h.** For each condition, statistical significance of prediction accuracies was determined in each subject’s native brain space and then projected into the MNI brain space. The number of subjects with significant prediction accuracy is shown for each voxel on the cortical surface of the MNI brain. The number of significant subjects is given by the color scale shown at bottom. Dark red voxels are significantly predicted in all subjects, and dark blue voxels are not significantly predicted in any subjects. **e.** In the Single Words condition, no voxels are semantically selective for any subjects. **f.** In the Semantic Blocks condition, scattered voxels in left STS are semantically selective in two out of four subjects. **g.** In the Sentences condition, voxels in the bilateral STS, STG, angular gyrus, ventral precuneus, and SFS are semantically selective in two out of four subjects. **h.** In the Narratives condition, voxels in bilateral angular gyrus, bilateral STS, anterior temporal lobe, SFS, SFG, IFG, ventral precuneus, posterior cingulate gyrus, and right STG are semantically selective in all four subjects. The results shown here are consistent with those in Figure 4, and they suggest that increasing context increases the representation of semantic information across the cortical surface at the group level but not for individual subjects.

Because the Narratives condition contains more contextual information than the other three conditions, we hypothesized that we would find more semantically selective voxels in the Narratives condition than in the other three conditions. To test this, we calculated the difference in the number of semantically selective voxels between the Narratives condition and each of the other three conditions. The difference between the Narratives and Single Words conditions is 4745, 7355, 7786, and 1757 voxels for subjects 1-4, respectively (p<0.05 for all subjects). The difference between the Narratives and Semantic Blocks conditions is 4037, 7355, 7786, and 1757 voxels for subjects 1-4, respectively (p<0.05 for all subjects). Finally, the difference between the Narratives and Sentences conditions is 3179, 4774, 7786, and 1757 voxels for subjects 1-4, respectively (p<0.05 for all subjects). The difference between the Narratives and Single Words conditions partly reflects the fact that most voxels have low evoked SNR in the Single Words condition and high evoked SNR in the Narratives condition (Figure 3). Because it is impossible to model noise, differences in evoked SNR across conditions directly affect the number of voxels that achieve a significant model fit. The difference between the Narratives and Semantic Blocks conditions also partly reflects differences in evoked SNR --for most voxels, evoked SNR is low in the Semantic Blocks condition and high for the Narratives condition (Figure 3). In contrast, the evoked SNR is high for many voxels in both the Narratives and the Sentences conditions (Figure 3), so the difference in the number of semantically selective voxels is unlikely to be due to differences in evoked SNR. Instead, this result suggests that semantic information is represented more widely across the cortical surface in the Narratives condition than in the Sentences condition.

To determine which semantic concepts are represented in voxels that are semantically selective in the Narratives condition but not in the Sentences condition, we looked at the semantic tuning of such voxels. The semantic tuning of each voxel is given by its 985-dimensional vector of estimated semantic model weights, one weight for each of the 985 semantic model features (see Methods). Since the semantic model has 985 features, it is difficult and impractical to interpret the semantic tuning of a voxel by looking at each individual semantic feature directly. Instead, we projected each voxel’s estimated semantic model weights into a low-dimensional subspace of the semantic model, and interpreted semantic tuning based on how the semantic weights projected into this subspace. This low-dimensional subspace was created by applying principal component analysis (PCA) to the aggregated estimated semantic model weights of seven subjects in Huth et al. 2016. Applying PCA to the aggregated semantic model weights returns principal components (PCs) that are ordered by how much variance they explain in the aggregated semantic model weights. The low-dimensional subspace was defined as the first three PCs of the aggregated semantic model weights.

To visualize semantic tuning, we projected the estimated Narratives semantic model weights for each voxel onto the three PCs, and then we colored each voxel with an RGB color scheme. For each voxel, the red value indicates the projection onto the first PC, the green value indicates the projection onto the second PC, and the blue value indicates the projection onto the third PC. Figure 6 shows the estimated Narratives semantic model weights projected onto the three PCs for two subjects (S1 and S2, this analysis was not performed for S3 and S4 because they did not have any semantically selective voxels in the Sentences condition). In both subjects, most voxels that are semantically selective in the Narratives condition but not in the Sentences condition have either a high red value or a high green value. A high red value corresponds to tuning for concepts related to humans and social relationships, and a high green value corresponds to tuning for concepts related to materials and measurements. Thus, voxels that are semantically selective in the Narratives condition but not in the Sentences condition are tuned to these two semantic categories.

**Figure 6.**
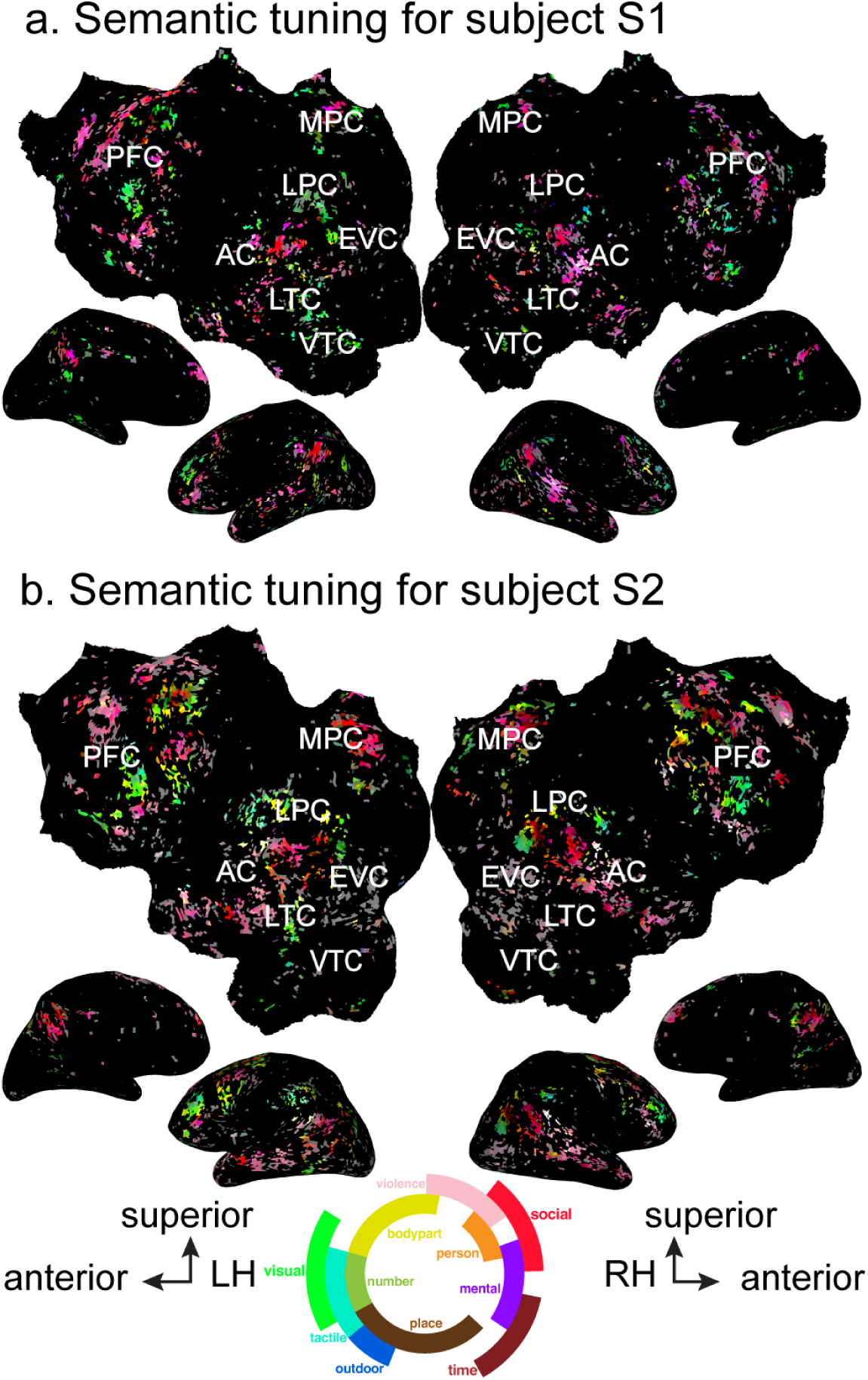
Semantic tuning of voxels that are semantically selective in the Narratives condition but not the Sentences condition. Semantic tuning is shown on the flattened cortical surface of two subjects (S1 and S2) for voxels that are semantically selective in the Narratives condition but not in the Sentences condition. These voxels are in the bilateral superior temporal sulcus, middle temporal gyrus, precuneus, inferior frontal gyrus, and ventrolateral and dorsolateral prefrontal cortex. Semantic model weights estimated in the Narratives condition were projected into a low-dimensional subspace created by performing principal components analysis (PCA) on semantic model weights estimated in Huth et al. 2016. Each voxel is colored according to the projection of its Narratives semantic model weights onto the first (red), second (green), and third (blue) PCs. The color wheel legend shows the semantic concepts associated with different colors. Most voxels in both subjects have a high red value or a high green value. A high red value corresponds to tuning for concepts related to humans and social relationships, and a high green value corresponds to tuning for concepts related to materials and measurements. (LH: Left Hemisphere, RH: Right Hemisphere, AC: auditory cortex, EVC: early visual cortex, LTC: lateral temporal cortex, VTC: ventral temporal cortex, LPC: lateral parietal cortex, MPC: medial parietal cortex, PFC: prefrontal cortex)

Differences in semantic representation between the Sentences and Narratives conditions could be limited to a difference in the number of voxels recruited to represent semantic information in each condition. However, we hypothesized that differences in contextual information between the two conditions could also lead to differences in semantic tuning in the voxels that are semantically selective in both conditions. To test this hypothesis, the semantic model weights estimated in the Sentences condition were correlated with the semantic model weights estimated in the Narratives condition for voxels that are semantically selective in both conditions. Figure 7 shows Pearson’s correlation coefficient between the semantic model weights estimated in the Sentences condition and the semantic model weights estimated in the Narratives condition mapped onto the cortical surface of two subjects (S1 and S2). In both subjects, semantic model weights for the Sentences and Narratives conditions are on average moderately correlated (S1 correlation min=-0.319, max=0.817, mean=0.344; S2 correlation min=-0.271, max=0.725, mean=0.316). This result shows that semantic tuning changes in semantically selective voxels between the Sentences and Narratives conditions.

**Figure 7.**
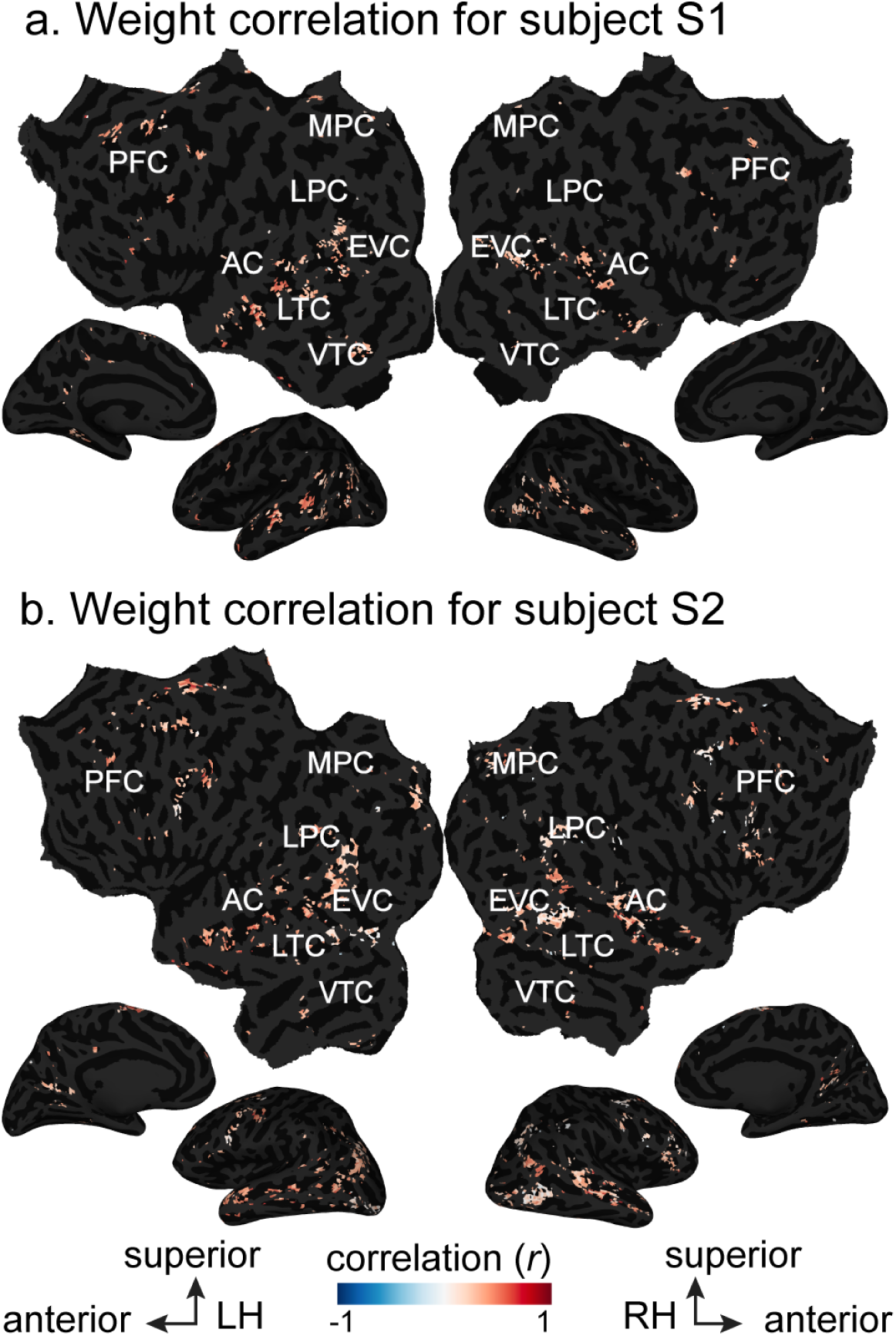
Correlation of semantic model weights estimated in the Sentences and Narratives conditions. Pearson’s correlation coefficient between semantic model weights estimated in the Sentences condition and semantic model weights estimated in the Narratives conditions is plotted on the flattened cortical surface of two subjects (S1 and S2). Only voxels that are semantically selective in both conditions are shown. These include voxels in the superior temporal sulcus and prefrontal cortex in both hemispheres and in both subjects. These voxels are on average moderately correlated between these two conditions (S1 correlation min=-0.319, max=0.817, mean=0.344; S2 correlation min=-0.271, max=0.725, mean=0.316), indicating that the semantic model weights estimated in the Sentences and Narratives conditions point in different directions in the semantic space. This shows that semantic tuning changes between the Sentences and Narratives conditions. (LH: Left Hemisphere, RH: Right Hemisphere, AC: auditory cortex, EVC: early visual cortex, LTC: lateral temporal cortex, VTC: ventral temporal cortex, LPC: lateral parietal cortex, MPC: medial parietal cortex, PFC: prefrontal cortex)

To determine how semantic tuning changes between the Sentences and Narratives conditions, we looked at how estimated semantic model weights differ between the two conditions. For every voxel that is semantically selective in both conditions, we subtracted its semantic model weights estimated in the Sentences condition from its semantic model weights estimated in the Narratives condition (see Methods). The resulting semantic difference vector describes the semantic concept that changes between the voxel’s semantic tuning in the Sentences and Narratives conditions. The difference vector resides in the same 985-dimensional semantic space as the semantic model weights, so we projected the difference vector into a low-dimensional semantic subspace to interpret its semantic tuning. This subspace was created by applying PCA to the difference vectors for each subject separately. The first five PCs explained 47.1% of the variance in subject S1 and 48.2% of the variance in subject S2 (see Extended Data Figure 8-1 for partial scree plots), indicating that the semantic tuning shifts can be described by a relatively low number of dimensions. Figure 8 shows the projection of the difference vectors onto the first three PCs for one subject (S1; see Extended Data Figure 8-2 for subject S2). Each voxel is colored according to how positively (red) or negatively (blue) its difference vector projects onto each of the three PCs. For the first PC, voxels in bilateral STS and bilateral SFG have a strong positive projection while voxels in bilateral angular gyrus have a strong negative projection in both subjects. For the second PC, voxels in bilateral angular gyrus and superior STS have a strong positive projection in both subjects. No voxels have a strong negative projection in either subject. For the third PC, voxels in right STS have a strong positive projection in both subjects. No voxels have a strong negative projection in either subject. These results suggest that semantic tuning shifts between the Sentences and Narratives conditions are spatially organized across cortex.

**Figure 8.**
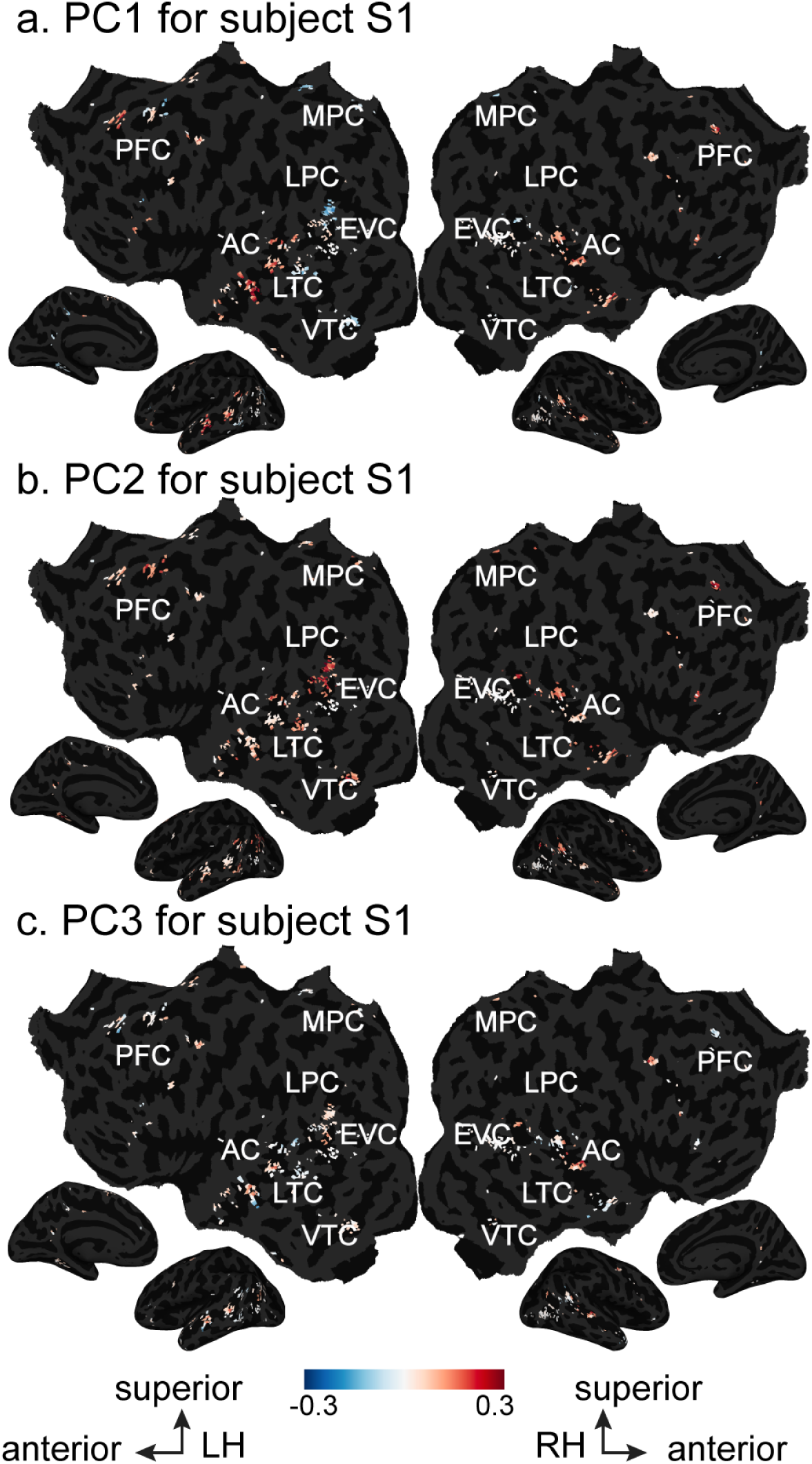
Semantic tuning shifts between the Sentences and Narratives conditions. Semantic model weights estimated in the Sentences condition were subtracted from semantic model weights estimated in the Narratives condition. PCA was then applied to the resulting difference vectors for each subject separately. The projection of the difference vectors onto the first three PCs is shown on the flattened cortical surface of one subject (S1; see Extended Data Figure 8-2 for subject S2; see Extended Data Figure 8-1 for the amount of variance explained by each of the first five PCs for each subject). Only voxels that are semantically selective in both conditions are shown. Projection strength is given by the color scales, and the ends of the color scales are labeled with the corresponding semantic concepts for each PC. Voxels that project onto one end of a PC appear red, while voxels that project onto the opposite end of the same PC appear blue. (LH: Left Hemisphere, RH: Right Hemisphere, AC: auditory cortex, EVC: early visual cortex, LTC: lateral temporal cortex, VTC: ventral temporal cortex, LPC: lateral parietal cortex, MPC: medial parietal cortex, PFC: prefrontal cortex) **a.** The first PC for subject S1 is shown. Voxels in bilateral STS and bilateral SFG are red while voxels in bilateral angular gyrus are blue in both subjects. **b.** The second PC for subject S1 is shown. Voxels in bilateral angular gyrus and superior STS are red while no voxels are blue in both subjects. **c.** The third PC for subject S1 is shown. Voxels in right STS are red while no voxels are blue in both subjects. The ten most and least correlated words for each PC are shown in Extended Data Figure 8-3. These results show that semantic tuning shifts between the Sentences and Narratives conditions are spatially organized across cortex.

To interpret the PCs of the semantic difference vectors, we looked at the words in the semantic model that were correlated with each PC (see Extended Data Figure 8-3 for the ten most correlated and least correlated words for each PC for each subject). For subject S1, the first PC is high on words related to interviewing and interrogation and low on words related to building and investing. The second PC is high on words related to packages and deliveries and low on words related to athletics. The third PC is high on words related to measurement and low on words related to family. For subject S2, the first PC is high on words related to visualization and low on words related to time and numbers. The second PC is high on words related to travel and deliveries and low on words related to body parts and actions. The third PC is high on function words and words related to numbers and low on informal words and interjections. The first three PCs for subject S1 are only moderately correlated to the first three PCs for subject S2: the correlation for the first PC is 0.3144, the correlation for the second PC is 0.5996, and the correlation for the third PC is 0.2351. This suggests that semantic tuning shifts between the Sentences and Narratives conditions are subject-dependent. However, additional analysis using a larger subject pool is needed to determine the individual differences in semantic tuning.

So far we have shown that semantic information is represented more widely across the cortical surface in the Narratives condition compared to the Single Words, Semantic Blocks or Sentences conditions (Figures 4, 5, 6). Next, we wanted to assess whether semantic model weights estimated using stimuli with little context can generalize to natural stimuli. Due to the low evoked SNR and low semantic model predictions in the Single Words and Semantic Blocks conditions (Figure 3) (Extended Data Figure 4-2), we hypothesized that the semantic model weights estimated in these conditions would generalize more poorly to the Narratives condition than the Sentences condition. To examine this, we used the semantic model weights estimated in the conditions with less context than Narratives (Single Words, Semantic Blocks, or Sentences) to predict brain activity in the Narratives condition. We then compared these cross-condition predictions to within-condition predictions (Narratives predicting Narratives condition). Figure 9 shows the results of this analysis in subject S1 (see Extended Data Figure 9-2 and Figure 9-3 for subject S2). Visual inspection of Figure 9 shows that when semantic model weights estimated in the Single Words condition are used to predict the Narratives condition only scattered voxels across the cerebral cortex are predicted (Figure 9a, blue voxels) but no voxel is predicted in the bilateral temporal, parietal, and prefrontal regions involved in within-condition predictions using Narratives (Figure 9a, red voxels). When semantic model weights estimated in the Semantic Blocks condition are used to predict the Narratives condition only scattered voxels across the cerebral cortex are predicted (Figure 9b, blue voxels). A few voxels in left STS are well predicted in cross-condition predictions (Semantic Blocks predicting Narratives) and in within-condition predictions using Narratives (Figure 9b, white voxels). Most of the remaining voxels within the bilateral temporal, parietal, and prefrontal regions are only well predicted in within-condition predictions using Narratives (Figure 9b, red voxels). In contrast, when semantic model weights estimated in the Sentences condition are used to predict the Narratives condition voxels in bilateral angular gyrus, bilateral STS, bilateral TPJ, bilateral sPMv, bilateral ventral precuneus, bilateral SFG, bilateral IFG, and left SFS are well predicted in cross-condition predictions (Figure 9c white voxels; See Extended Data Figure 9-1 for significance, p<0.05, FDR corrected). These voxels are also well predicted in within-condition predictions using Narratives (Figure 9c, white voxels). In addition, there are voxels within the bilateral temporal, parietal, and prefrontal regions that are only well predicted in within-condition predictions using Narratives. These voxels are located in left IPL, right SFS, bilateral STG, right anterior temporal lobe, and bilateral posterior cingulate gyrus (Figure 9c, red voxels). Scattered voxels located in bilateral precuneus, right IFG, and portions of SFS are not well predicted in within-condition predictions using Narratives but are well predicted in cross-condition predictions (Sentences predicting Narratives) (Figure 9c, blue voxels). These results show that stimuli with little context (Single Words or Semantic Blocks) do not generalize well to stimuli that has more context than isolated single words (Sentences or Narratives). In addition, estimated model weights using Sentences generalize well in some voxels within the temporal, parietal, and prefrontal regions. However, remaining voxels in these regions are only well predicted by a semantic model that is trained on natural stories stimuli (Narratives) than single isolated sentences (Sentences).

**Figure 9.**
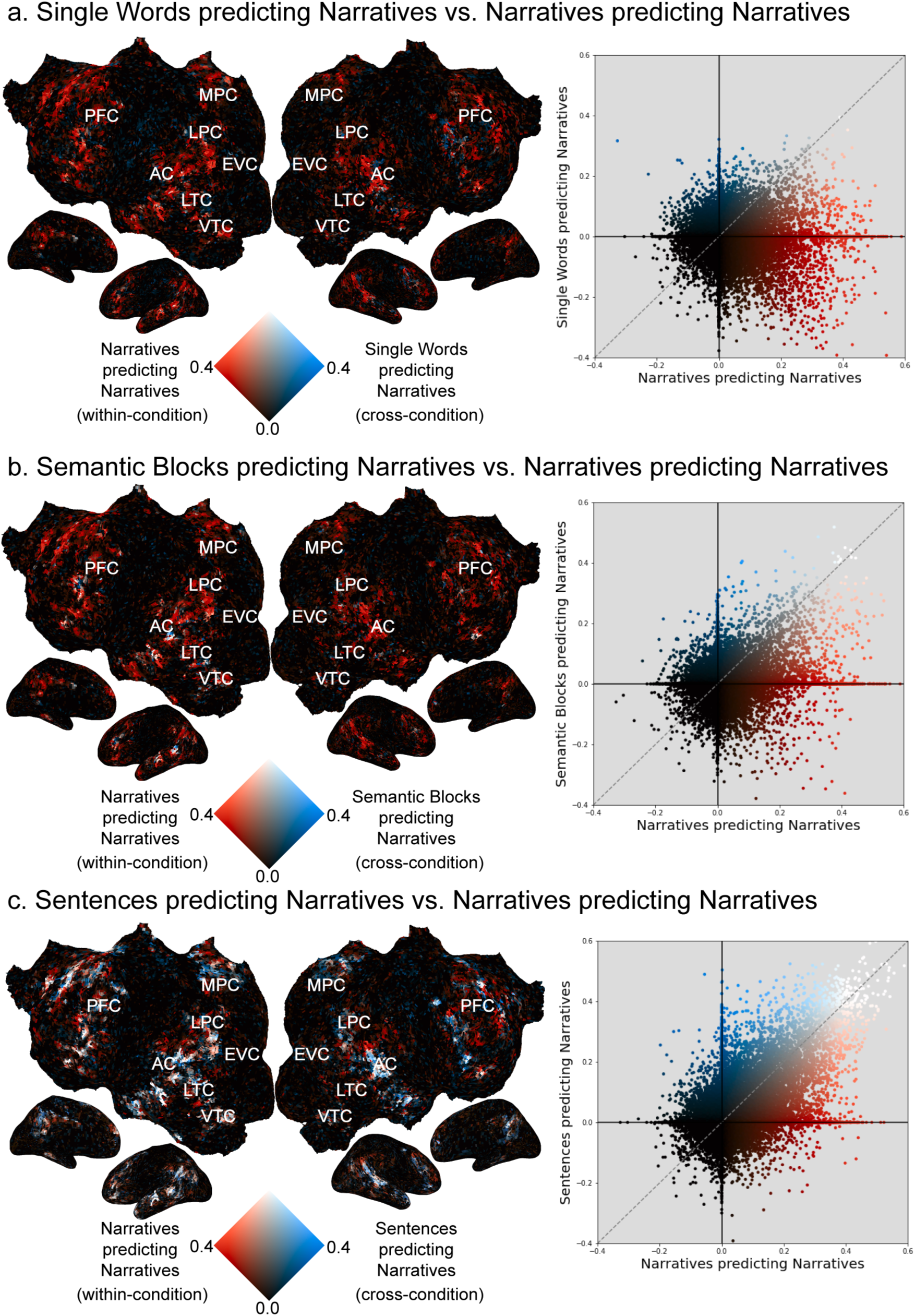
Generalization of semantic model weights estimated in the Single Words, Semantic Blocks, and Sentences conditions to the Narratives condition for subject S1. **a.** Semantic model weights estimated in the Single Words condition were used to predict BOLD responses to the held-out validation stimulus in the Narratives condition. (left) The resulting cross-condition semantic model prediction accuracies are shown with the within-condition Narratives semantic model prediction accuracies on the flattened cortical surface of subject S1 with a 2D colormap (see Extended Data Figure 9-2 for subject S2). (LH: Left Hemisphere, RH: Right Hemisphere, AC: auditory cortex, EVC: early visual cortex, LTC: lateral temporal cortex, VTC: ventral temporal cortex, LPC: lateral parietal cortex, MPC: medial parietal cortex, PFC: prefrontal cortex) The axes of the colormap correspond to the cross-condition (blue) and within-condition (red) prediction accuracies. Voxels where the within-condition prediction accuracy is high and the cross-condition prediction accuracy is low appear red. Voxels where the within-condition prediction accuracy is low and the cross-condition prediction accuracy is high appear blue. Voxels where both the within-condition prediction accuracy and the cross-condition prediction accuracy are high appear white. Finally, voxels where both the within-condition prediction accuracy and the cross-condition prediction accuracy are low appear black. In this comparison, many voxels throughout bilateral temporal, parietal, and prefrontal cortex are red. In addition, there are a few blue and white voxels scattered across the cortical surface. (right) Cross-condition semantic model prediction accuracy (y-axis) is plotted against within-condition Narratives semantic model prediction accuracy (x-axis) for each cortical voxel. In most voxels, the cross-condition prediction accuracy is worse than the Narratives prediction accuracy. **b.** Semantic model weights estimated in the Semantic Blocks condition were used to predict BOLD responses to the held-out validation stimulus in the Narratives condition. The format is the same as panel a. Many voxels across bilateral temporal, parietal, and prefrontal cortex are red. Voxels located in the left superior temporal sulcus (STS) are white, and a few voxels scattered across the cortical surface are blue. In most voxels, the cross-condition prediction accuracy is worse than the Narratives prediction accuracy. **c.** Semantic model weights estimated in the Sentences condition were used to predict BOLD responses to the held-out validation stimulus in the Narratives condition. The format is the same as panel a. Voxels located in left IPL, right SFS, bilateral STG and bilateral posterior cingulate gyrus are red. Voxels located in bilateral angular gyrus, bilateral STS, portions of TPJ, in bilateral sPMv, bilateral ventral precuneus, bilateral SFG, bilateral IFG, and left SFS are white. These cross-condition prediction accuracy in these white voxels also reach statistical significance. This suggests that semantic model weights estimated in the Sentences condition generalize to the Narratives condition in these voxels (see Extended Data Figure 9-1 for S1 and Extended Data Figure 9-3 for S2). Scattered voxels located in bilateral precuneus, right IFG, and portions of SFS are blue. In many voxels, the cross-condition prediction accuracy is worse than the Narratives prediction accuracy. Together, these results show semantic model weights estimated in conditions with less context do not generalize well to natural stories.

## Discussion

The aim of this study was to determine whether and how context affects semantic representations in the human brain. Our results show that both evoked SNR and semantic representations are affected by the amount of context in the stimulus. First, stimuli with relatively more context (Narratives, Sentences) evoke brain responses with higher SNR compared to stimuli with relatively less context (Semantic Blocks, Single Words) (Figure 3). Second, increasing the amount of context increases the representation of semantic information across the cortical surface at the group level (Figures 4, 5). However, in individual subjects, only the Narratives condition consistently increased the representation of semantic information compared to the Single Words condition (Figures 4, 5). Third, increasing the amount of context changes the semantic tuning of semantically selective voxels across the cortical surface (Figures 6, 7, 8). These results strongly imply that neuroimaging studies that use isolated words or sentences do not fully map the functional brain representations that underlie natural language comprehension (Figure 9). By using the voxelwise encoding modeling approach with a specific semantic feature space, we demonstrate for the first time where semantic information is represented when different levels of contextual information are present in the stimuli. Thus, our results are much more specific to semantic representations than results in past studies.

Our observations that increasing context increases both the evoked SNR and the cortical representation of semantic information at the group level are fully consistent with results from previous neuroimaging studies. Several previous studies found that stimuli with more context evoke larger, more widespread patterns of brain activity (Mazoyer et al. 1993; Xu et al. 2005; Jobard et al. 2007), that brain activity evoked for individual words is modulated by context (Just, Wang, and Cherkassky 2017), and that brain activity evoked by stimuli with more context are more reliable than those evoked by stimuli with less context (Lerner et al. 2011). Furthermore, previous studies that used narrative stimuli (Wehbe et al. 2014; Huth et al. 2016; Pereira et al. 2018; Deniz et al. 2019; Hsu et al. 2019; Popham et al. 2021) identified many more voxels involved in semantic processing than studies that used isolated words or sentences (for reviews see (Jeffrey R. Binder et al. 2009; Price 2010, 2012).

Our results are also consistent with prior studies that have shown a broadly distributed semantic network that represents the meaning of language (Huth et al. 2016; Jeffrey R. Binder et al. 2009; Popham et al. 2021). One of the interesting aspects of the semantic network is that each semantic concept appears to be represented in multiple distinct brain areas. One potential hypothesis is that these repeated patterns actually represent different aspects of each of the semantic concepts, but they appear to be the same because of current limitations in our ability to measure and model brain activity. If this is true, then one might expect that selectivity in this network would increase as a subject focuses on a concept for a longer period of time, or as increasing semantic context is provided.

However, there are several important differences between the results we reported here and those reported in previous neuroimaging studies. First, past studies that used isolated sentences found left IFG involved in semantic processing (Constable et al. 2004; Rodd, Davis, and Johnsrude 2005; Humphries et al. 2007). We found few semantically selective voxels scattered in left IFG in two out of four subjects in the Sentences condition (Figures 4 and 5). Second, past studies that used isolated words found bilateral STS, bilateral lateral sulcus, left IFG, left MTG, and left ITG involved in lexical processing (Mazoyer et al. 1993; Booth et al. 2002; Xu et al. 2005; Jobard et al. 2007; Lerner et al. 2011). In contrast, we did not find any semantically selective voxels in the Single Words condition (Figures 4 and 5). Finally, one previous study looked at brain activity evoked by a stimulus conceptually similar to Semantic Blocks (Mollica et al. 2020). In the study, Mollica et al. (2020) used sentences that were scrambled such that nearby words could be combined into meaningful phrases. They found that the brain activity evoked by scrambled sentences was similar to the brain activity evoked by unscrambled sentences in left IFG, left middle frontal gyrus, left temporal lobe, and left angular gyrus. In contrast, we found voxels that were semantically selective in both the Semantic Blocks and Sentences conditions in left STS (Figures 4 and 5). We only found a few scattered voxels in IFG that were semantically selective in both of these conditions, and this result was not consistent across subjects (Figures 4-2 and 5). However, we also found that voxels in IFG were well predicted when the semantic model estimated in the Sentences condition was used to predict data in the Narratives condition (Figure 9 and Extended Data Figure 9-1). These two results suggest that IFG was involved in semantic processing when subjects read sentences (e.g. Sentences and Narratives conditions) and not when subjects read words that were semantically in context but were shown in no particular word order to the subjects (e.g. in the Semantic Blocks condition).

The inconsistencies between this study and past studies most likely stem from five major methodological differences between this study and those earlier studies. First, we avoided smoothing our data before performing analyses. We performed our analyses for each subject in their native brain space, and we did not perform any spatial smoothing across voxels. In contrast, most previous studies performed normalization procedures to transform their data into a standard brain space and applied a spatial smoothing operation across voxels (Lindquist 2008; Carp 2012). Spatial smoothing and normalization procedures can incorrectly assign signal to voxels and average away meaningful signal and individual variability in language processing (Steinmetz and Seitz 1991; Fedorenko and Kanwisher 2009; Fedorenko, Duncan, and Kanwisher 2012; Huth et al. 2016; Deniz et al. 2019). Thus, brain regions identified by past studies may be more relevant at the group level than in individual subjects. These smoothing procedures likely contribute to the inconsistencies observed between past studies and this study.

Second, we used an explicit computational model to identify semantically selective voxels. In contrast, most previous studies identified semantic brain regions by contrasting different experimental conditions (Jeffrey R. Binder et al. 2008, 2009; Price 2012). Although past studies designed their experimental conditions to isolate brain activity involved in semantic processing (Jeffrey R. Binder et al. 2008, 2009), there could be unexpected differences unrelated to semantic processing between the conditions. For example, experiments that contrast a semantic task with a phonological task (Jeffrey R. Binder et al. 2008, 2009) may have task difficulty as a confound. As a result, it is possible that some semantic brain areas identified by past studies are actually involved in processing the unexpected differences rather than semantics. We would likely not have identified such brain areas in this study, since our semantic model only contains information about semantics.

Third, we evaluated semantic model prediction accuracy on a separate, held-out validation dataset. In contrast, most previous studies drew inferences from analyses performed on only one dataset without a validation dataset (Jeffrey R. Binder et al. 2009). Performing analyses on only one dataset can lead to inflated results that are overfit to the dataset (Soch, Haynes, and Allefeld 2016). Thus, some semantic brain areas identified by past studies may only be relevant for the specific stimuli, experimental design, or data used in those studies. Such study-specific brain areas would not generalize to other studies, such as this study.

Fourth, we collected a relatively large amount of fMRI data per subject from four subjects. In contrast, most previous studies collected a small amount of fMRI data per subject from many (15-30) subjects. Because fMRI data is noisy, most previous studies either averaged their data across subjects and/or smoothed their data to observe the effects of interest. However, as discussed earlier, smoothing and averaging fMRI data can lead to incomplete conclusions about language processing in the brain (Steinmetz and Seitz, 1991; Fedorenko and Kanwisher, 2009; Fedorenko et al., 2012; Huth et al., 2016; Deniz et al., 2019). In this study, we avoided averaging across subjects and smoothing procedures by collecting a relatively large amount of data per subject. Moreover, each subject provided a complete replication of all analyses because each subject had their own model fitting and validation data. Thus, even though there are fewer subjects in this study than in previous studies, it is likely that our findings will generalize to new subjects.

Finally, subjects in our study passively read the stimulus words, which allowed us to directly compare the Narratives condition with the other three conditions. In contrast, many past studies of semantic processing used active tasks involving lexical decisions (J. R. Binder et al. 2003), matching (Vandenberghe et al. 1996), or monitoring (Démonet et al. 1992). Active tasks are thought to increase subject engagement, which can increase evoked BOLD SNR. Thus, if we had used an active task, the effect of context on evoked SNR might have been even larger than the differences that we report here. In addition, different active tasks can affect semantic processing differently in the brain (Toneva et al. 2020). Therefore, task effects likely contributed to the inconsistencies observed between past studies and this study.

To our knowledge, no previous language neuroimaging studies have looked at whether stimulus context affects semantic tuning. One interesting aspect of our results is that the semantic tuning shifts are different for subjects S1 and S2. One potential explanation for the discrepancy across subjects could be noise. Another possible explanation is that since both subjects saw the same stimuli in the Sentences and Narratives conditions, the difference in tuning shifts could be due to individual differences in attention rather than differences in the stimuli. In the absence of narrative structure subjects likely attend to different parts in a sentence, whereas when there is narrative structure subjects likely attend to similar semantic categories lead by the general narrative arc. This explanation is consistent with a previous study from our lab showing that many voxels across cortex shift their tuning towards attended semantic categories (Çukur et al. 2013). However, further research needs to be conducted about context-dependent semantic tuning shifts during language comprehension.

Many language neuroimaging studies use isolated sentences to localize the language network (e.g., (Fedorenko et al. 2010; Scott, Gallée, and Fedorenko 2017; Wilson et al. 2017)). These localizers contrast isolated sentences with non-words (i.e., sentences > non-words) to identify regions of interest (ROIs) in the brain involved in language processing. The identified ROIs often include left IFG, left middle frontal gyrus, left temporal lobe, left angular gyrus, and right temporal lobe. Consistent with these localizers, many voxels in the listed ROIs have high EV in the Sentences condition. In fact, the raw EV value in the Sentences condition is higher than the raw EV value in the Narratives condition in many voxels, suggesting that the Sentences condition engages the language network more than the Narratives condition. More predictable stimuli could lead to less activation the second time the subject reads the stimuli that would lead to lower EV. In addition, we find fewer semantically selective voxels in the Sentences condition than in the Narratives condition in all subjects (Figure 4 and Figure 5). Instead, we find that out of the five feature spaces we used in this study, the “number of letters” feature space has the highest prediction accuracy in the Sentences condition in all subjects. This suggests that a substantial portion of brain activations evoked by isolated sentences reflects the effect of low level features. However, the variance in the Sentences condition could also be explained by a different feature space that we did not include in our analyses for this paper.

Our study used a semantic model to determine whether and how semantic representations change across the four conditions. Although our semantic model is able to capture the semantic properties of individual words, it nonetheless has some limitations. First, because this model likely captures some narrative information that is correlated with word-level semantic information, some of the brain activity predicted by our semantic model may therefore reflect higher-level linguistic or domain-general representations (Fedorenko, Duncan, and Kanwisher 2012; Blank and Fedorenko 2017). Second, our semantic model has one static embedding for each word, and it does not differentiate between different word senses or different contexts in which a word may appear. Therefore, our semantic model may not predict voxel activity as well as other models that integrate contextual semantic information differently (Toneva and Wehbe 2019; Jain and Huth 2018; Heilbron et al. 2022; Schmitt et al. 2021; Goldstein et al. 2022; Schrimpf et al. 2021), specifically in the Sentences and Narratives conditions. The voxelwise modeling framework provides a straightforward method for evaluating alternative semantic models directly by construction of appropriate feature spaces. Therefore, a valuable direction for future research would be to examine other semantic models, and to include language models that explicitly account for factors such as contextual information, narrative structure, metaphor, and humor.

In conclusion, our results show that increasing the amount of stimulus context increases the SNR of evoked brain responses, increases the representation of semantic information in the brain, and affects the semantic tuning of semantically selective voxels. These results imply that neuroimaging studies that use isolated words or sentences to study semantic processing or to localize the language network (Fedorenko et al. 2010) may provide a misleading picture of semantic language comprehension in daily life. Although natural language stimuli are much more complex than isolated words and sentences, the development and validation of the voxelwise encoding model framework for language processing (Huth et al. 2016; de Heer et al. 2017; Deniz et al. 2019; Popham et al. 2021) has made it possible to rigorously test hypotheses about semantic processing with natural language stimuli. To ensure that the results of neuroimaging study generalize to natural language processing, we suggest that future studies of semantic processing should use more naturalistic stimuli.

---

**Figure 3-1.**
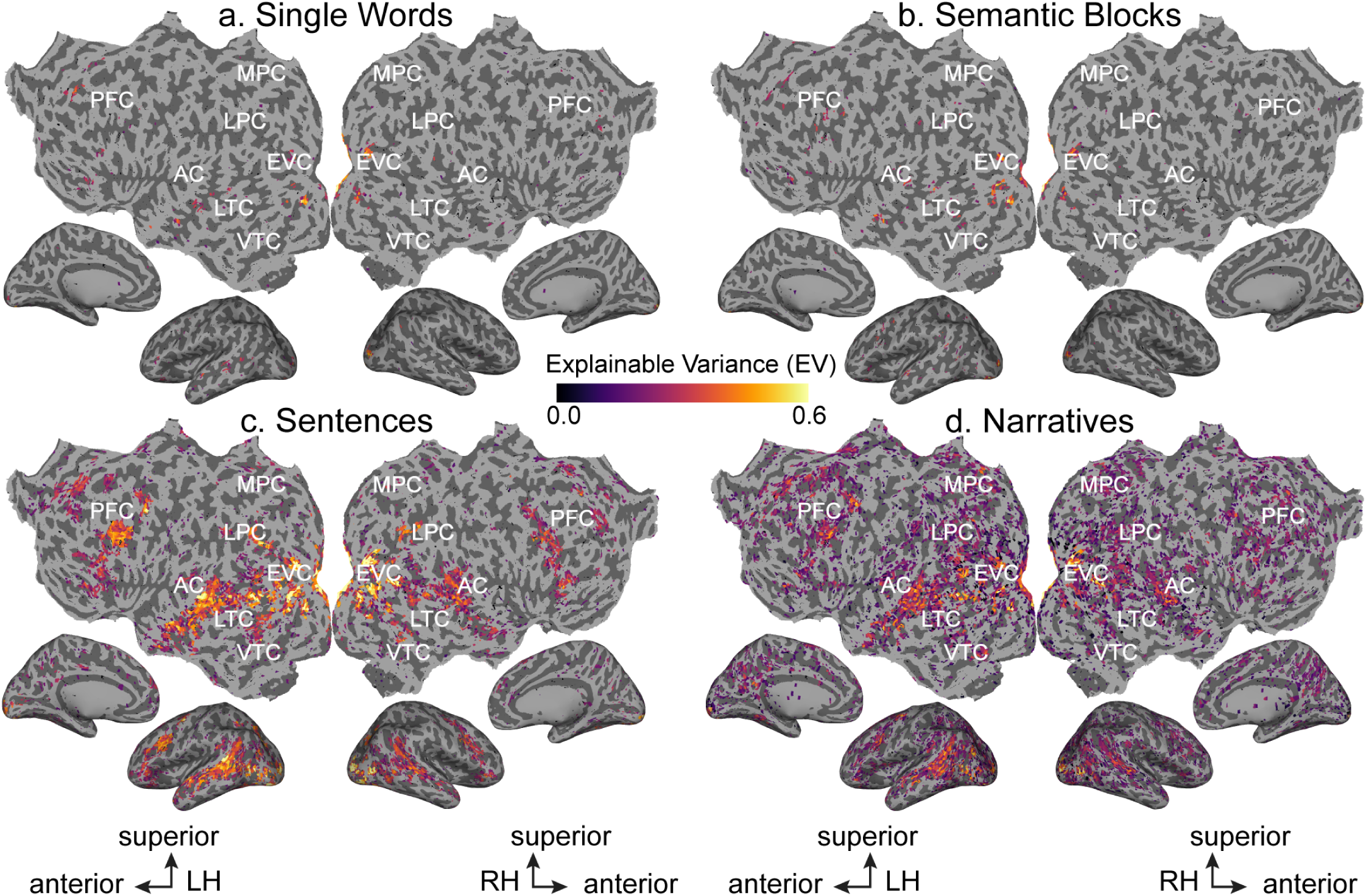
Significant explainable variance (EV) for the four conditions across the cortical surface. EV is shown for the four conditions on the flattened cortical surface of one subject (S1). EV was computed as an estimate of the evoked signal-to-noise ratio (SNR). Only voxels with significant EV (p<0.05, FDR corrected) are shown. EV is given by the color scale shown in the middle, and voxels that have high EV appear yellow. Voxels with EV values that are not statistically significant are shown in gray. (LH: Left Hemisphere, RH: Right Hemisphere, AC: auditory cortex, EVC: early visual cortex, LTC: lateral temporal cortex, VTC: ventral temporal cortex, LPC: lateral parietal cortex, MPC: medial parietal cortex, PFC: prefrontal cortex) **a.** EV was computed for the Single Words condition, and significant voxels are shown on the flattened cortical surface of subject S1. Scattered voxels in bilateral primary visual cortex, left STS, and left IFG have significant EV. **b.** Same as panel **a**. but for the Semantic Blocks condition. Similar to the Single Words condition, scattered voxels in bilateral primary visual cortex, left STS, and left IFG have significant EV. **c.** Same as panel **a**. but for the Sentences condition. Many voxels in bilateral visual, parietal, temporal, and prefrontal cortices have significant EV. **d.** Same as panel **a**. but for the Narratives condition. Similar to the Sentences condition, voxels in bilateral visual, parietal, temporal, and prefrontal cortices have high EV.

**Figure 3-2.**
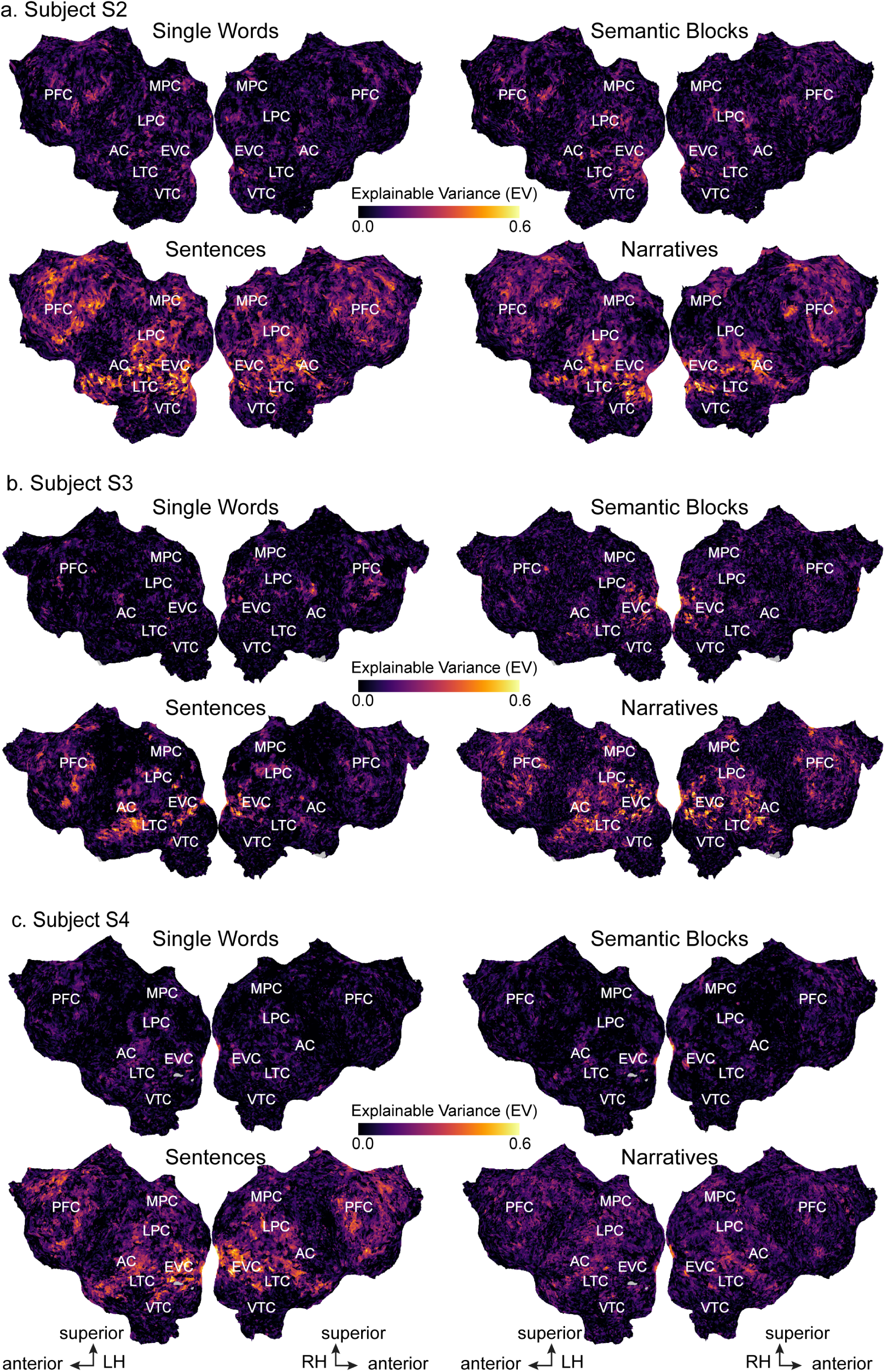
Explainable variance (EV) for the four conditions across the cortical surface for subjects S2, S3, and S4. EV is shown for the four conditions on the flattened cortical surface of subjects S2, S3, and S4. The format is the same as Figure 3. EV was computed as an estimate of the evoked signal-to-noise ratio (SNR). EV is given by the color scale shown in the middle, and voxels that have high EV (i.e., high evoked SNR) appear yellow. (LH: Left Hemisphere, RH: Right Hemisphere) Across all subjects, EV is low across most of the cortical surface in the Single Words and Semantic Blocks conditions. In contrast, EV is high for many voxels in bilateral visual, parietal, temporal, and prefrontal cortices in the Sentences and Narratives conditions.

**Figure 4-1.**
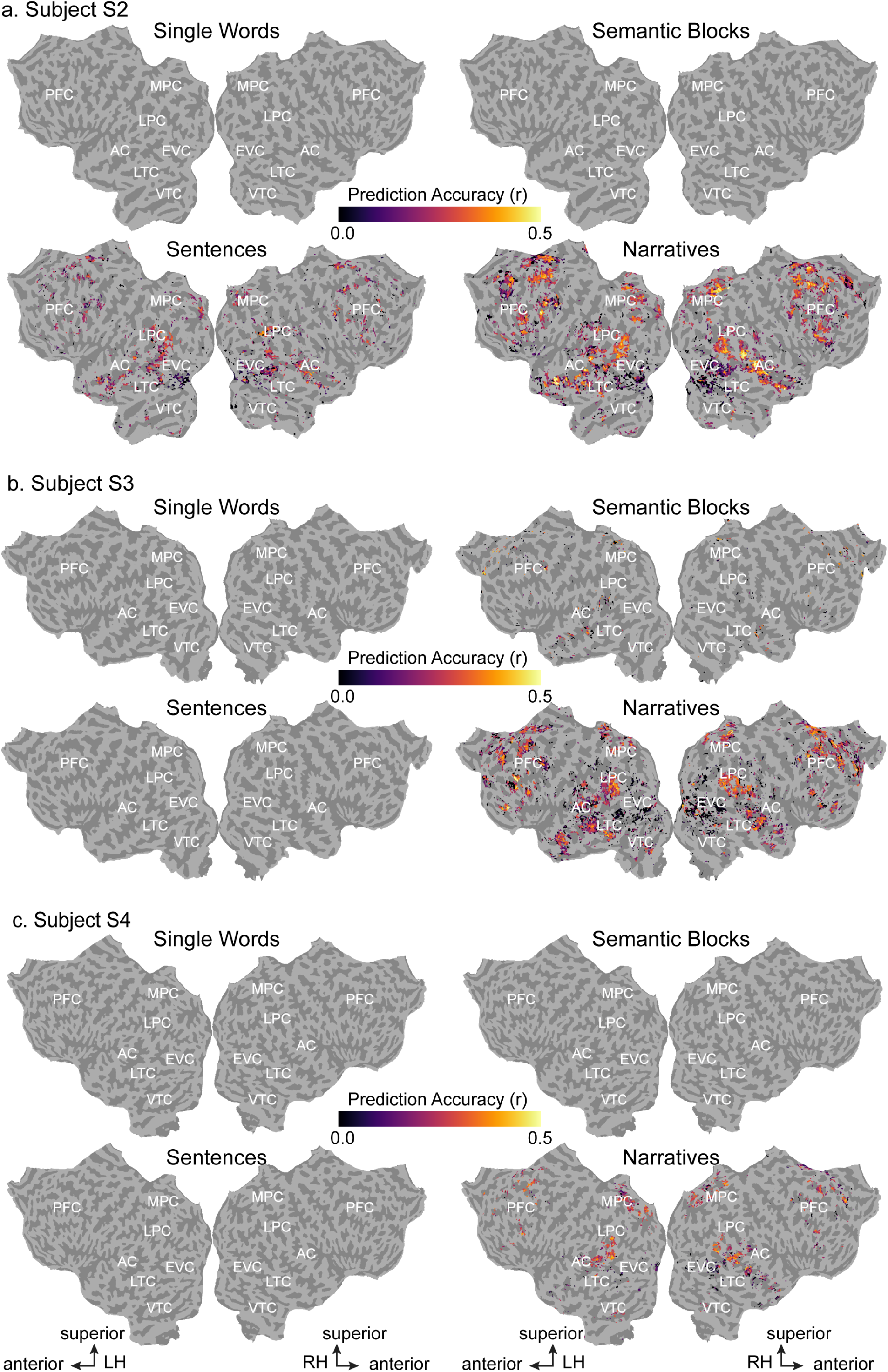
Semantic model prediction accuracy for the four conditions across the cortical surface for subjects S2, S3, and S4. Semantic model prediction accuracy in the four conditions is shown on the flattened cortical surface of subjects S2, S3 and S4. The format is the same as Figure 4. Voxelwise modeling was first used to estimate semantic model weights in the four conditions. Semantic model prediction accuracy was then computed as the correlation (r) between the subject’s recorded BOLD activity to the held-out validation story and the BOLD activity predicted by the semantic model. In each panel, only voxels with significant semantic model prediction accuracy (p<0.05, FDR corrected) are shown. Prediction accuracy is given by the color scale in the middle, and voxels that have a high prediction accuracy appear yellow. Voxels with semantic model prediction accuracies that are not statistically significant are shown in gray. (LH: Left Hemisphere, RH: Right Hemisphere) In the Single Words condition, no voxels are significantly predicted in all subjects. In the Semantic Blocks condition, scattered voxels in left STS, left angular gyrus, left sPMv, and bilateral SFS are significantly predicted in subject S3. In the Sentences condition, voxels in bilateral STS, bilateral STG, bilateral angular gyrus, bilateral ventral precuneus, bilateral SFS and SFG, bilateral IFG, and bilateral sPMv are significantly predicted in subject S2. In the Narratives condition, voxels in bilateral angular gyrus, bilateral ventral precuneus, bilateral SFS and SFG, and right STS are significantly predicted in all three subjects. In addition, bilateral STG, left STS, bilateral Broca’s area and IFG, and bilateral sPMv are significantly predicted in subjects S2 and S3.

**Figure 4-2.**
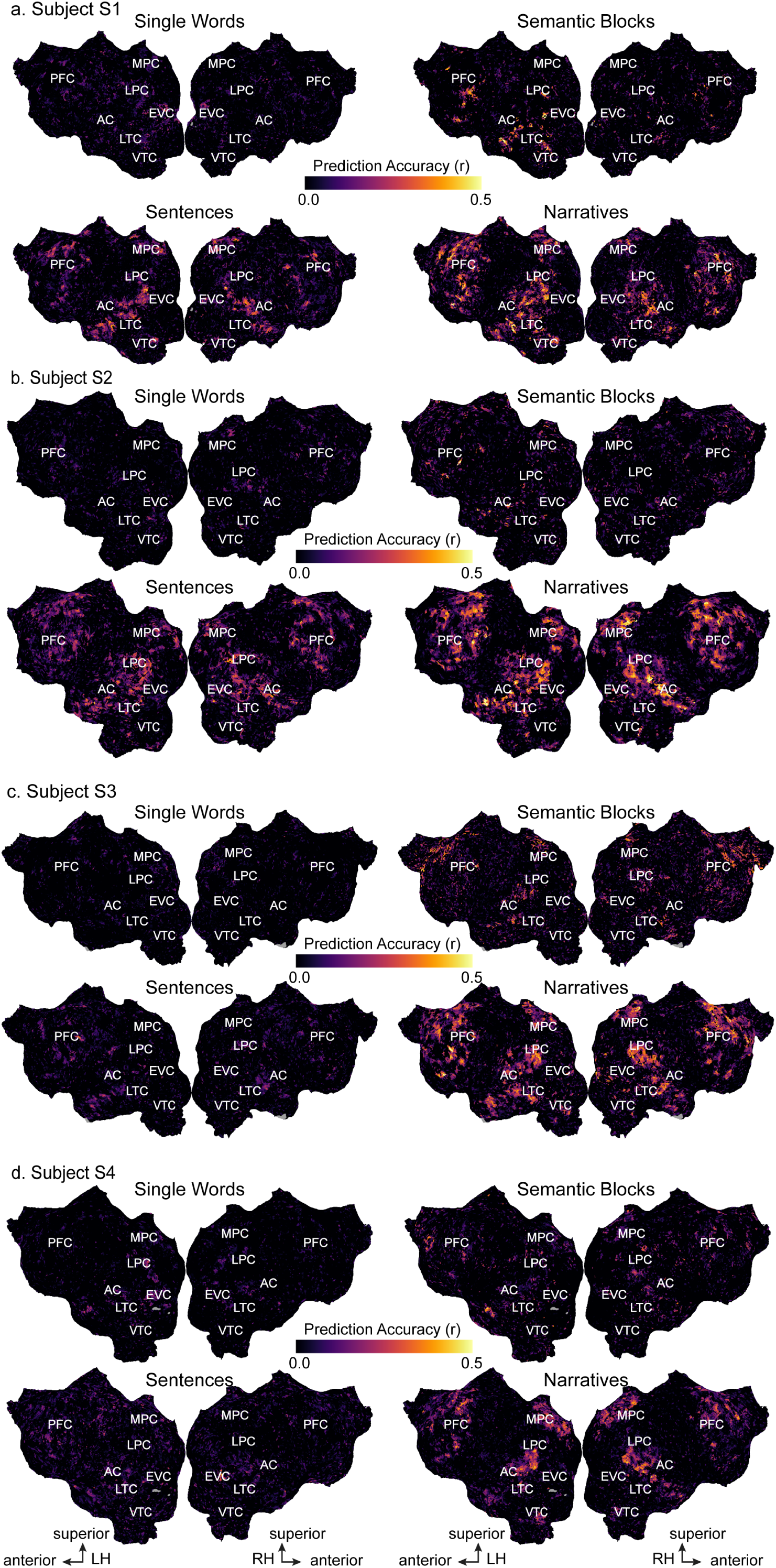
Un-thresholded semantic model prediction accuracy for the four conditions across the cortical surface for all subjects. Un-thresholded semantic model prediction accuracy in the four conditions is shown for all subjects on each subject’s flattened cortical surface. Voxelwise modeling was first used to estimate semantic model weights in the four conditions. Semantic model prediction accuracy was then computed as the correlation (r) between the subject’s recorded BOLD activity to the held-out validation story and the BOLD activity predicted by the semantic model. Prediction accuracy is given by the color scale in the middle, and voxels that have a high prediction accuracy appear yellow. (LH: Left Hemisphere, RH: Right Hemisphere, AC: auditory cortex, EVC: early visual cortex, LTC: lateral temporal cortex, VTC: ventral temporal cortex, LPC: lateral parietal cortex, MPC: medial parietal cortex, PFC: prefrontal cortex) In the Single Words condition, prediction accuracy is high in scattered voxels in primary visual cortex in subjects S1 and S4. In the Semantic Blocks condition, prediction accuracy is high in voxels in left STS and left angular gyrus in subjects S1 and S3. In addition, prediction accuracy is high in voxels in left Broca’s area and IFG in subject S1, and prediction accuracy is high in voxels in bilateral SFS, SFG, and ventral precuneus in subject S3. In the Sentences condition, prediction accuracy is high in voxels in bilateral angular gyrus, STS, STG, MTG, anterior temporal lobe, IFG, sPMv, SFS, SFG, and ventral precuneus in subjects S1 and S2. In the Narratives condition, prediction accuracy is high in voxels in bilateral angular gyrus, STS, STG, MTG, anterior temporal lobe, Broca’s area and IFG, sPMv, SFS, SFG, ventral precuneus, and posterior cingulate gyrus in all subjects.

**Figure 4-3.**
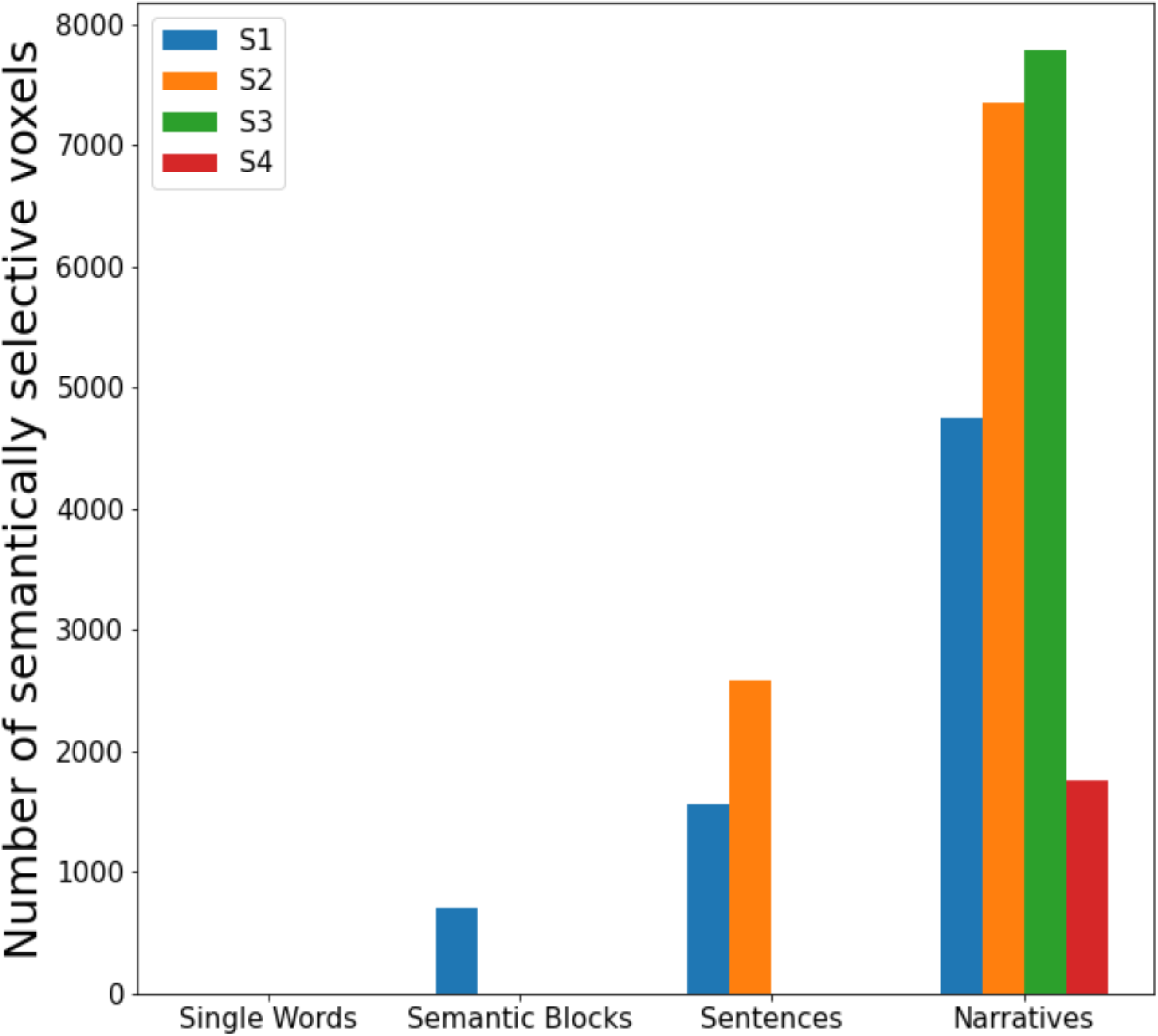
Number of semantically selective voxels for the four conditions for all subjects. The number of semantically selective voxels for each subject is plotted for the four conditions. In the Single Words condition, no voxels are semantically selective in any of the four subjects. In the Semantic Blocks condition, the number of semantically selective voxels is 708, 0, 0, and 0 for subjects 1-4, respectively. In the Sentences condition, the number of semantically selective voxels is 1566, 2581, 0, and 0 for subjects 1-4, respectively. In the Narratives condition, the number of semantically selective voxels is 4745, 7355, 7786, and 1757 for subjects 1-4, respectively.

**Figure 8-1.**
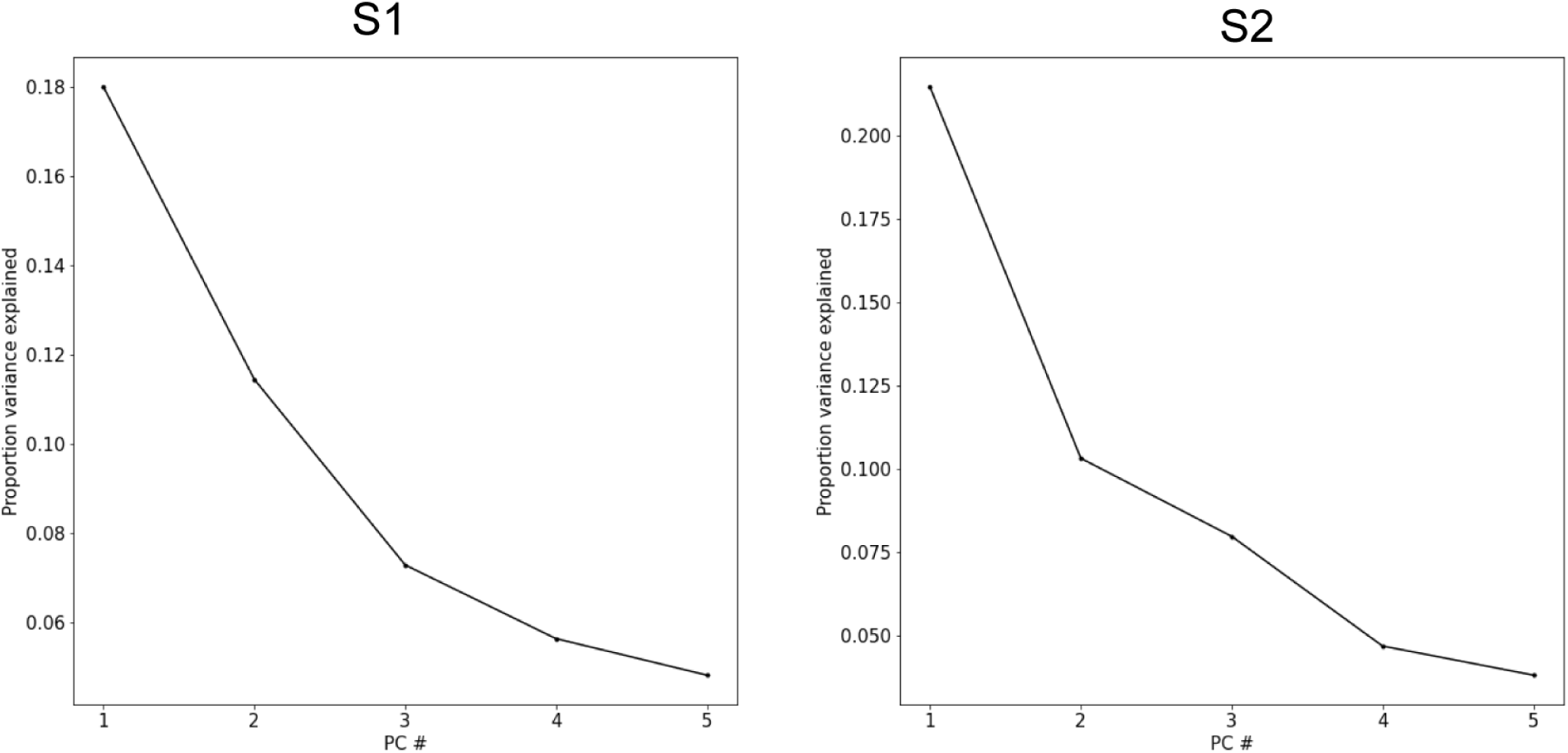
Proportion of variance explained by PCs of semantic difference vectors. Semantic difference vectors were computed by subtracting semantic model weights estimated in the Sentences condition from semantic model weights estimated in the Narratives condition. PCA was then applied to the difference vectors for each subject separately. The amount of variance explained by each of the first five PCs is plotted for each subject. The first five PCs explain 47.1% of the variance in subject S1 and 48.2% of the variance in subject S2.

**Figure 8-2.**
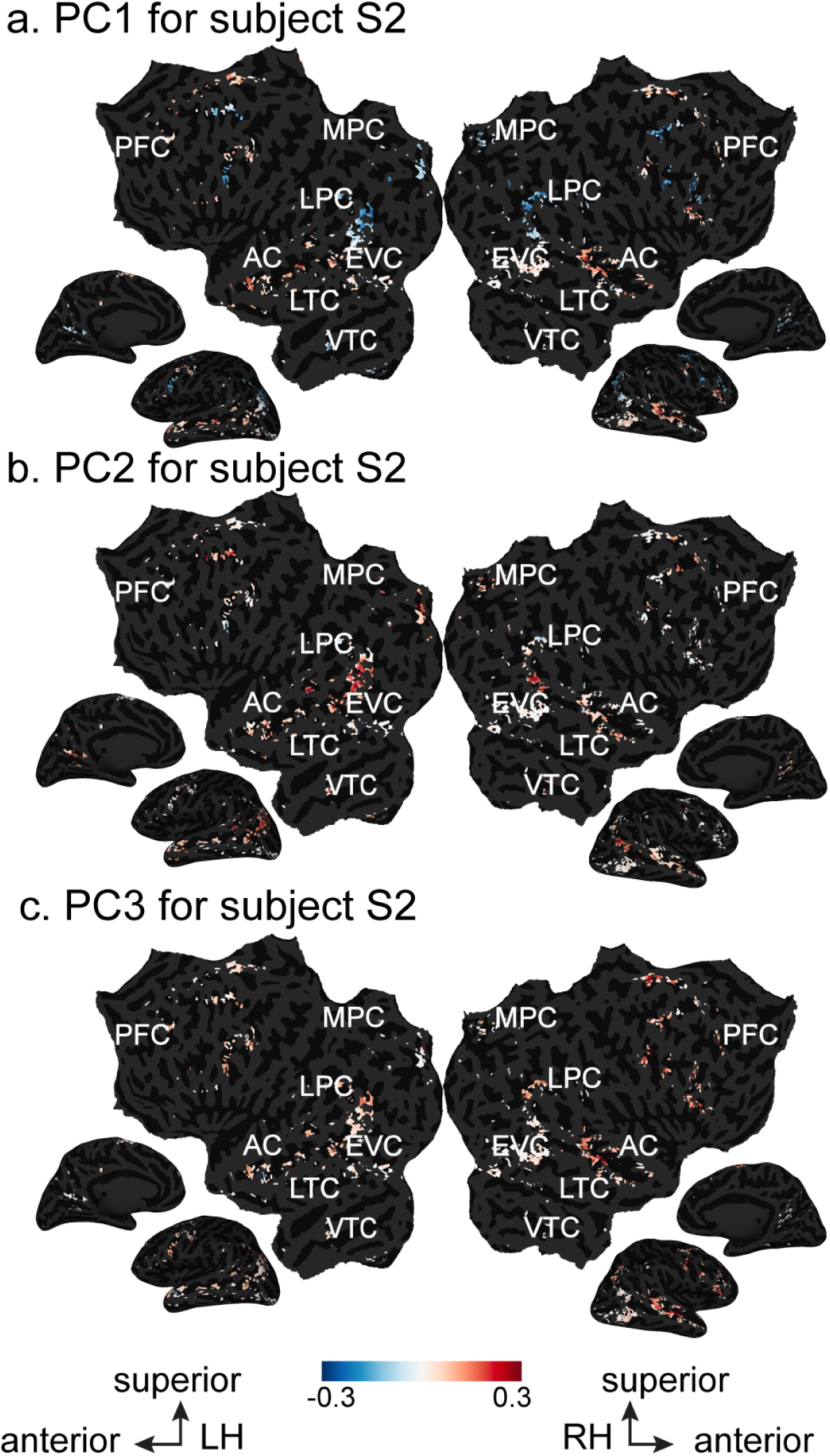
Semantic tuning shifts between the Sentences and Narratives conditions for subject S2. Semantic model weights estimated in the Sentences condition were subtracted from semantic model weights estimated in the Narratives condition. PCA was then applied to the resulting difference vectors for each subject separately. The projection of the difference vectors onto the first three PCs is shown on the flattened cortical surface of subject S2. Only voxels that are semantically selective in both conditions are shown. Projection value is given by the color scale in the middle. Voxels that project positively onto a PC appear red, while voxels that project negatively onto a PC appear blue. (LH: Left Hemisphere, RH: Right Hemisphere, AC: auditory cortex, EVC: early visual cortex, LTC: lateral temporal cortex, VTC: ventral temporal cortex, LPC: lateral parietal cortex, MPC: medial parietal cortex, PFC: prefrontal cortex) **a.** For the first PC, voxels in bilateral STS and bilateral SFG have a strong positive projection while voxels in bilateral angular gyrus, bilateral RSC, bilateral IFG, and bilateral SFS have a strong negative projection. **b.** For the second PC, voxels in bilateral angular gyrus, bilateral superior STS, bilateral RSC, and bilateral SFS have a strong positive projection while no voxels have a strong negative projection. **c.** For the third PC, voxels in right STS, bilateral angular gyrus, right SFS, and right IFG have a strong positive projection while no voxels have a strong negative projection.

**Figure 8-3.**
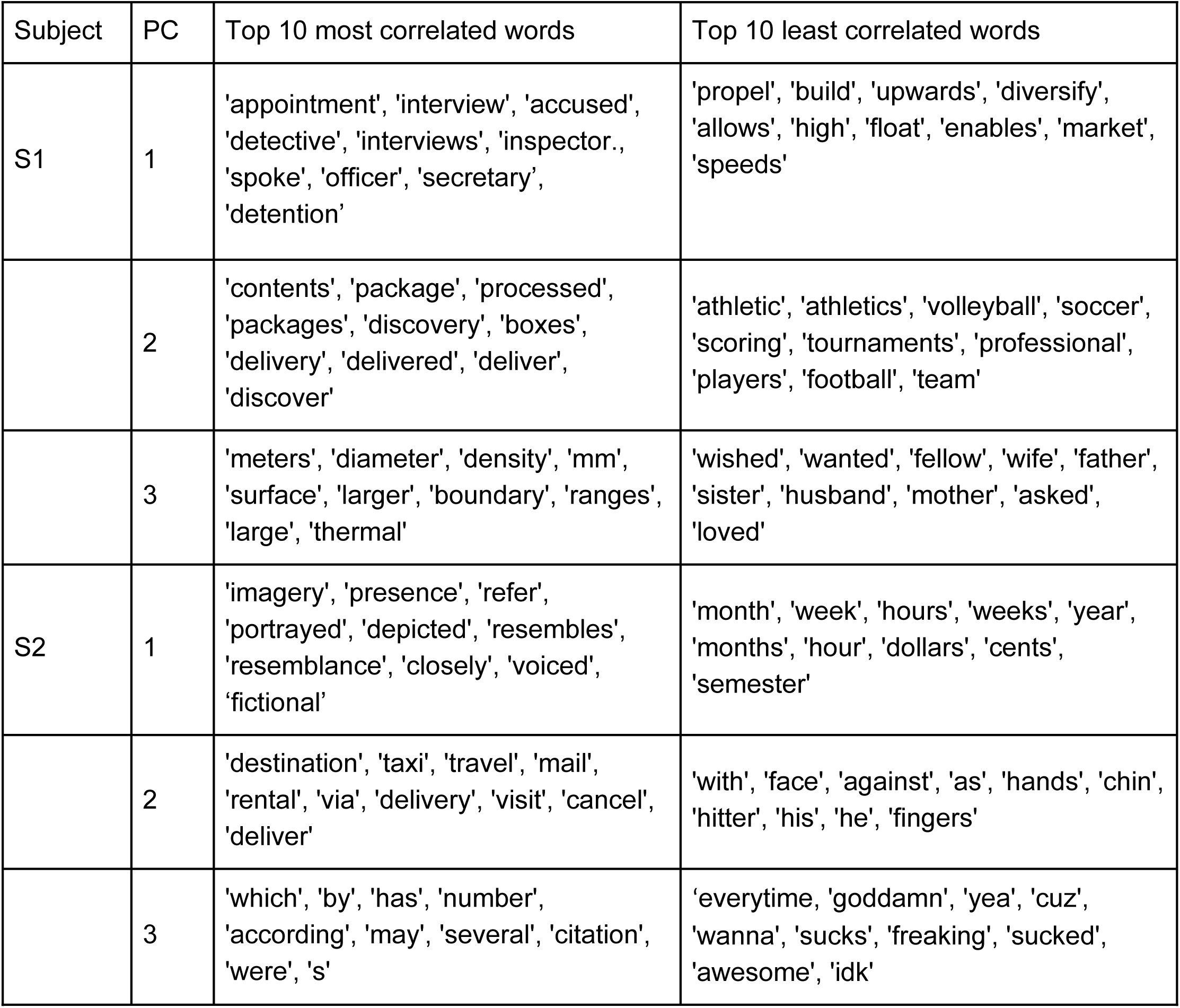
Most and least correlated words for each PC. The first three PCs of the difference vectors were correlated with words in the semantic model. The ten most correlated words and the ten least correlated words are shown for each PC for each subject.

**Figure 9-1.**
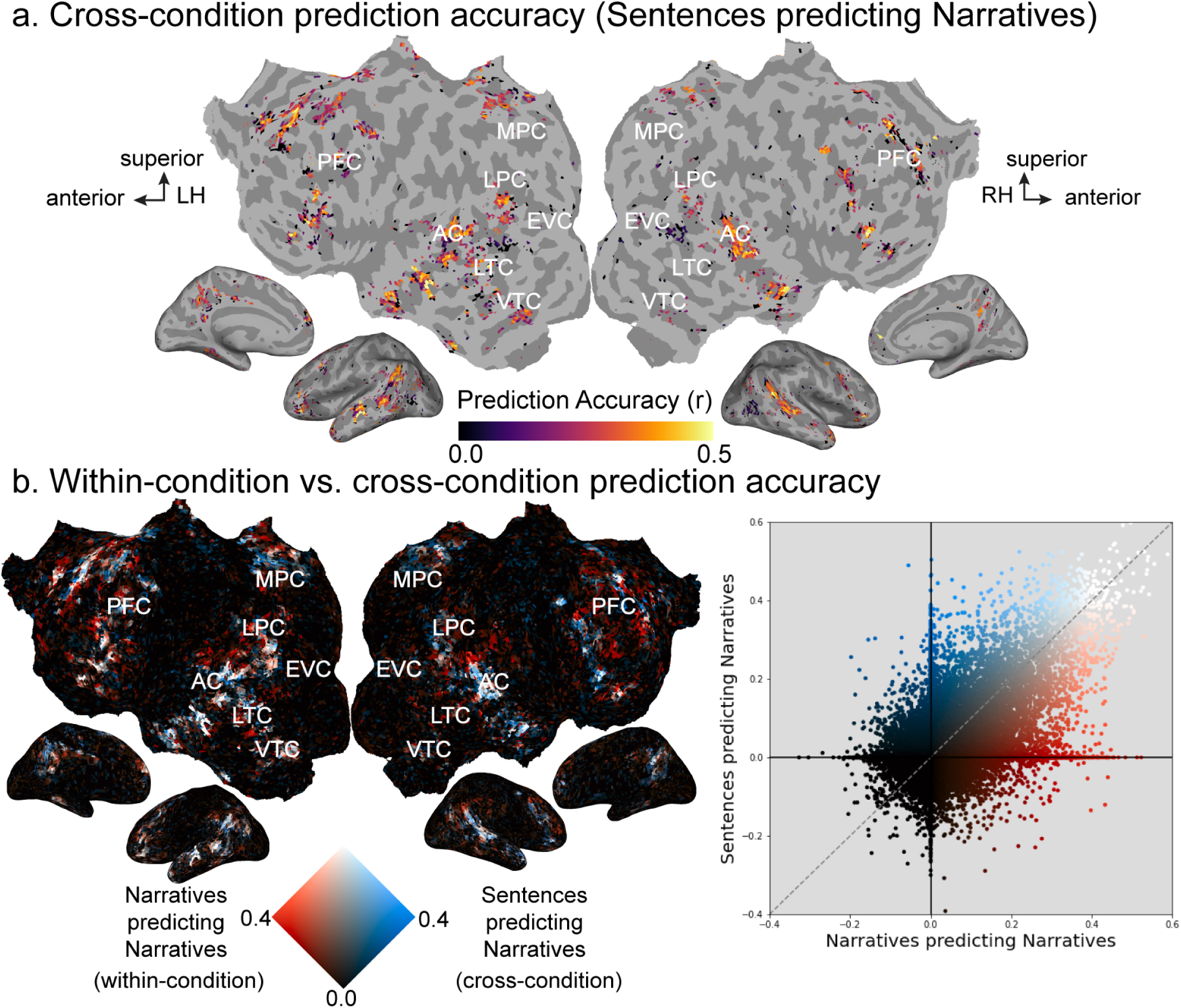
Cross-condition semantic model prediction accuracy for the Sentences and Narratives conditions. **a.** Semantic model weights estimated in the Sentences condition were used to predict BOLD responses to the held-out validation stimulus in the Narratives condition. The resulting cross-condition semantic model prediction accuracy is shown on the flattened cortical surface of one subject (S1; see Extended Data Figure 9-2 for S2). Only voxels with significant prediction accuracy (p<0.05, FDR corrected) are shown. Prediction accuracy is given by the color scale in the middle, and voxels that have a high prediction accuracy appear yellow. Voxels for which the cross-condition semantic model prediction accuracy is not statistically significant are shown in gray. (LH: Left Hemisphere, RH: Right Hemisphere, AC: auditory cortex, EVC: early visual cortex, LTC: lateral temporal cortex, VTC: ventral temporal cortex, LPC: lateral parietal cortex, MPC: medial parietal cortex, PFC: prefrontal cortex) Voxels in bilateral angular gyrus, bilateral STS, portions of TPJ, bilateral sPMv, bilateral ventral precuneus, bilateral SFG, bilateral IFG, and left SFS are significantly predicted. Semantic model weights estimated in the Sentences condition generalize to the Narratives condition in these voxels. **b.** Same panel as in Figure 9c depicted here for a direct comparison.

**Figure 9-2.**
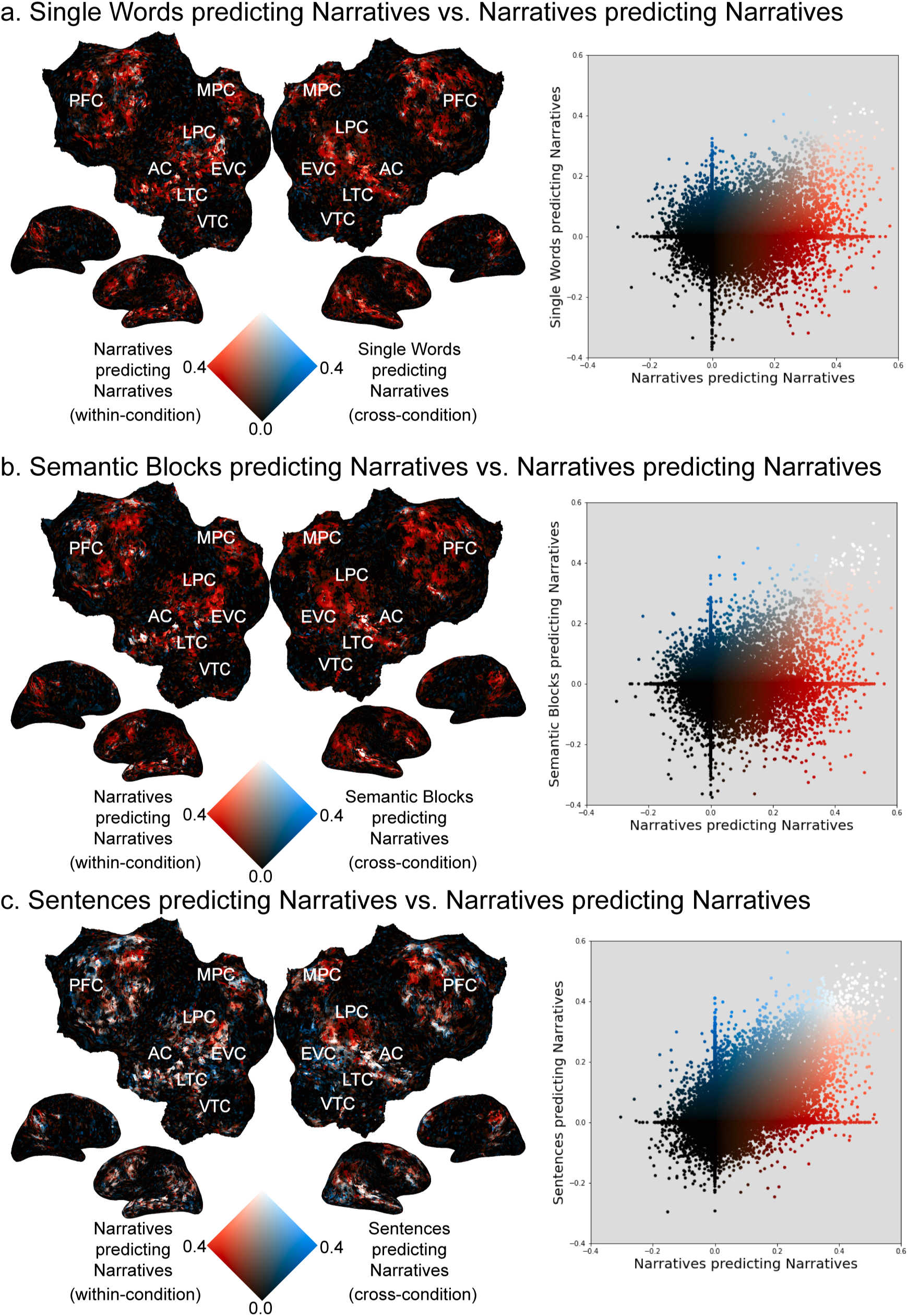
Generalization of semantic model weights estimated in the Single Words, Semantic Blocks, and Sentences conditions to the Narratives condition for subject S2. **a.** Semantic model weights estimated in the Single Words condition were used to predict BOLD responses to the held-out validation stimulus in the Narratives condition. (left) The resulting cross-condition semantic model prediction accuracies are shown with the within-condition Narratives semantic model prediction accuracies on the flattened cortical surface of subject S2 with a 2D colormap. (LH: Left Hemisphere, RH: Right Hemisphere, AC: auditory cortex, EVC: early visual cortex, LTC: lateral temporal cortex, VTC: ventral temporal cortex, LPC: lateral parietal cortex, MPC: medial parietal cortex, PFC: prefrontal cortex) The axes of the colormap correspond to the cross-condition (blue) and within-condition (red) prediction accuracies. Voxels where the within-condition prediction accuracy is high and the cross-condition prediction accuracy is low appear red. Voxels where the within-condition prediction accuracy is low and the cross-condition prediction accuracy is high appear blue. Voxels where both the within-condition prediction accuracy and the cross-condition prediction accuracy are high appear white. Finally, voxels where both the within-condition prediction accuracy and the cross-condition prediction accuracy are low appear black. In this comparison, many voxels throughout bilateral temporal, parietal, and prefrontal cortex are red. In addition, there are a few blue and white voxels scattered across the cortical surface. (right) Cross-condition semantic model prediction accuracy (y-axis) is plotted against within-condition Narratives semantic model prediction accuracy (x-axis) for each cortical voxel. In most voxels, the cross-condition prediction accuracy is worse than the Narratives prediction accuracy. **b.** Semantic model weights estimated in the Semantic Blocks condition were used to predict BOLD responses to the held-out validation stimulus in the Narratives condition. The format is the same as panel a. Many voxels across bilateral temporal, parietal, and prefrontal cortex are red. A few voxels located in the left superior temporal sulcus (STS) are white, and a few voxels scattered across the cortical surface are blue. In most voxels, the cross-condition prediction accuracy is worse than the Narratives prediction accuracy. **c.** Semantic model weights estimated in the Sentences condition were used to predict BOLD responses to the held-out validation stimulus in the Narratives condition. The format is the same as panel a. Voxels located in left IPL, right SFS and bilateral STG are red. Voxels located in bilateral angular gyrus, bilateral STS, portions of TPJ, in bilateral sPMv, bilateral SFG, bilateral IFG, and left SFS are white. These cross-condition prediction accuracy in these white voxels also reach statistical significance. This suggests that semantic model weights estimated in the Sentences condition generalize to the Narratives condition in these voxels (See Extended Data Figure 9-3). Scattered voxels located in bilateral precuneus, right IFG, and portions of SFS are blue. In many voxels, the cross-condition prediction accuracy is worse than the Narratives prediction accuracy. Together, these results show semantic model weights estimated in conditions with less context do not generalize well to natural stories.

**Figure 9-3.**
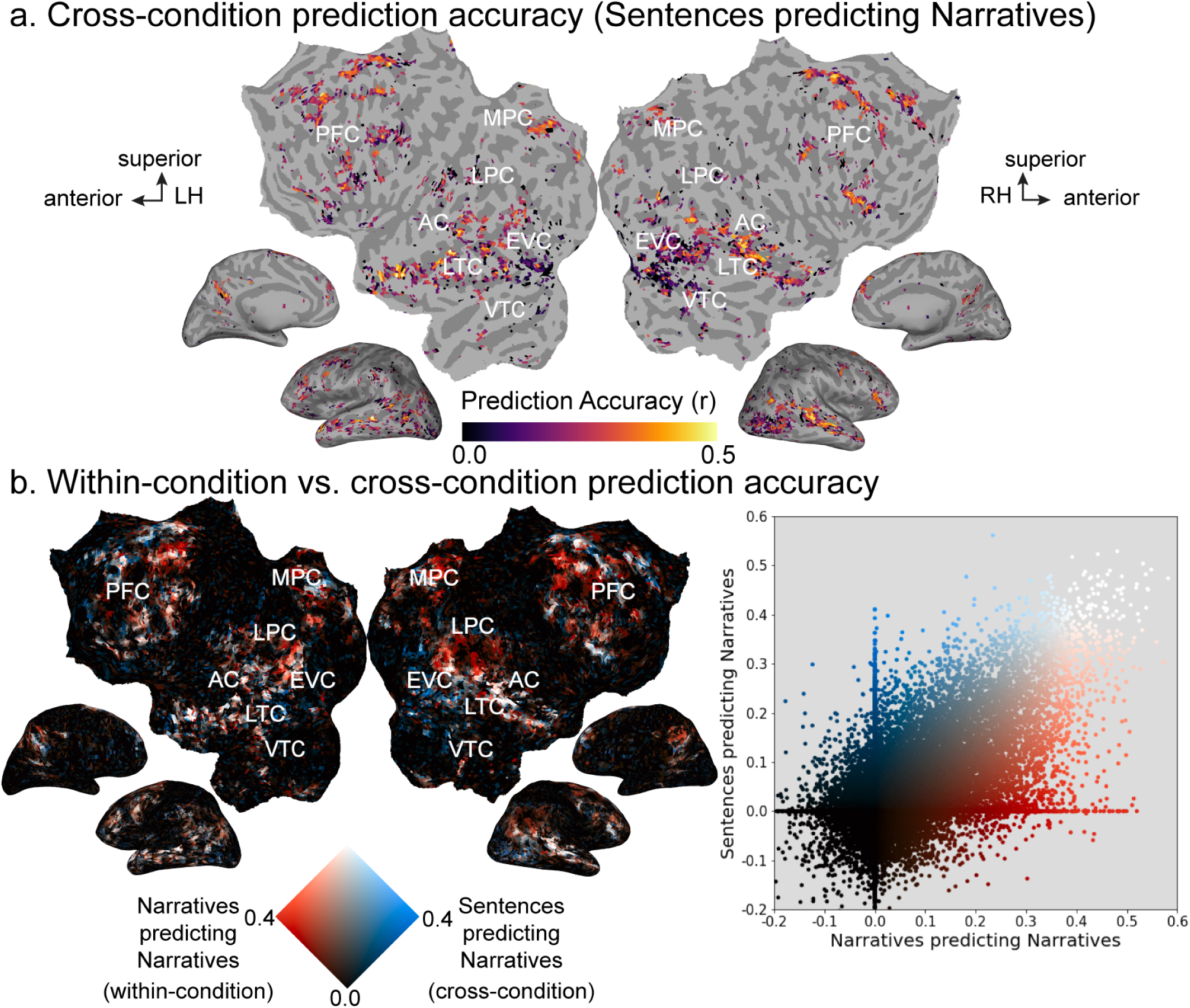
Prediction accuracy of semantic model weights estimated in the Sentences condition predicting data in the Narratives condition for subject S2. **a.** Semantic model weights estimated in the Sentences condition were used to predict BOLD responses for the held-out validation stimulus in the Narratives condition (cross-condition predictions). The resulting cross-condition semantic model prediction accuracy is shown on the flattened cortical surface of subject S2. Only voxels with significant prediction accuracy are shown (p<0.05, FDR corrected). Prediction accuracy is given by the color scale in the middle, and voxels that have a high prediction accuracy appear yellow. Voxels for which the cross-condition semantic model prediction accuracy is not statistically significant are shown in gray. (LH: Left Hemisphere, RH: Right Hemisphere, AC: auditory cortex, EVC: early visual cortex, LTC: lateral temporal cortex, VTC: ventral temporal cortex, LPC: lateral parietal cortex, MPC: medial parietal cortex, PFC: prefrontal cortex) Some voxels in bilateral angular gyrus, bilateral STS, portions of TPJ, in bilateral sPMv, bilateral ventral precuneus, bilateral SFG, bilateral IFG, and left SFS are significantly predicted when estimated semantic model weights in the Sentences condition are used to predict brain responses in the Narratives condition (p<0.05, FDR corrected). Sentences condition generalize well to Narratives condition in these voxels. **b.** Same panel as in Extended Data Figure 9-2c depicted here for a direct comparison.

